# Leveraging AI-powered interactive playbacks to decipher rules of communication in zebra finches

**DOI:** 10.64898/2026.02.12.705387

**Authors:** Logan S. James, Benjamin Hoffman, Jen-Yu Liu, Marius Miron, Milad Alizadeh, Emmanuel Fernandez, Matthieu Geist, Diane Kim, Aza Raskin, Jon T. Sakata, Emmanuel Chemla, Olivier Pietquin, Sarah C. Woolley

**Author notes:** These authors contributed equally to this work (Logan S. James, Benjamin Hoffman, Jen-Yu Liu). These authors listed alphabetically (Milad Alizadeh, Emmanuel Fernandez, Matthieu Geist, Diane Kim, Aza Raskin). Co-supervising authors (Emmanuel Chemla, Olivier Pietquin, Sarah C. Woolley).

## Abstract

Vocal interactions are fundamental for social functioning across animals, including humans. The diverse rules underlying these exchanges remain largely unknown, and emerging AI technologies offer promising avenues for investigation. We used computational tools to collect and analyze *>*1,000 hours of vocal interactions between female zebra finches and discovered that their interactions were characterized by correlated call production and structure, rapid acoustic modulation, and response selectivity. To test these interaction rules, we developed a generative audio large language model (ZF-AIM Acoustic Interaction Model) that engaged in real-time vocal exchanges with birds. When birds interacted with ZF-AIM, their vocal production and flexibility recapitulated key naturalistic features, which did not happen with non-interactive playbacks. Targeted ablations of ZF-AIM revealed that call timing and structure differentially contribute to natural vocal interactions. Using these AI-animal interactions, we demonstrate how AI can be leveraged to reveal fundamental rules underlying animal communication.

## Main text

Communicative exchanges in which individuals alternate signaling (turn-taking) are common in animals, including humans (*1–4*). During such exchanges, interlocutors often modulate the signals they produce based on the dynamics of the interaction; for instance, humans cue the end of a phrase with elongated words (*5, 6*), match the linguistic style of their interlocutor (*7*), and adjust response timing based on social connection and semantic content (*8*). Other animals also alternate signaling (*4, 9–13*) but the extent to which flexibility in timing and acoustic structures similarly underlie natural exchanges in non-human animals remains poorly understood. Specifically, experiments aimed at understanding vocal interactions in non-human animals are typically observational or reliant on brief playback experiments that are often static and lack naturalistic contingencies. Extended experiments using interactive playbacks that can be manipulated in targeted ways provide opportunities for understanding naturalistic vocal exchanges.

Recent developments in AI technologies (including large language models; LLMs) that take advantage of large annotated datasets of conversation have been used to not only model and simulate natural human conversation but also to reveal the impact of specific features (e.g., alignment (*14*), turn-taking (*15*), conversation length (*16*), persuasiveness (*17, 18*), and politeness (*19*)) on conversation outcomes. Moreover, LLM responses to questionnaires can approximate human responses and predict outcomes of psychological studies, suggesting they could complement or even replace human participants in some contexts (*20, 21*). Anticipating the potential applications of these technologies in the study of animal communication, we implemented AI-based analysis pipelines to decipher fundamental aspects of naturalistic vocal exchanges in finches, and developed an audio-LLM to engage in a sustained, realistic vocal interaction with an animal partner. We compared this interactive model to passive (non-interactive) playbacks and manipulated its behavior to reveal which factors are important for naturalistic conversations in zebra finches.

## Results

### 1. Female zebra finches exhibit dynamic flexibility in their vocal exchanges

Female zebra finches readily interact vocally using relatively simple calls, and these calls can impact group/breeding cohesion, individual recognition, and social bonding (*22–25*). Zebra finches rapidly respond to calls from other individuals (*26, 27*), and the timing of these responses is controlled by brain regions analogous to cortical sensorimotor areas underlying speech in humans (*28–30*). Despite how common these vocalizations are, little is known about how female finches use acoustic variation in their calls throughout an interaction.

We housed female zebra finches within a sound-attenuating chamber in cages separated by an opaque barrier, allowing acoustic but not visual communication (*31*), and analyzed > 830,000 calls from 40 pairwise interactions (24.5 hours of analysis per pair; n = 20 birds; Fig. 1A,B,C). We found clear evidence of vocal interactivity. Females produced bursts of calls separated by minutes of silence and the numbers of calls within these bursts was significantly correlated between calling partners (Fig. 1D,E; mixed-effects model: *p* < 0.001; Table S7). The number of calls also increased over time (*p* < 0.001). Birds produced very rapid responses to their partners’ calls (< 300 ms; compared to slower self repetitions; Fig. 1F), and we focused our further analyses on these responses. Specifically, we considered the first call initiated within 1 second following the end of a partner’s call to be a “direct response” (Figs. 1F, 7). While we used a relatively permissive threshold to classify direct responses, we found similar effects with both shorter and longer thresholds (e.g., nearly identical results with a 0.5 and 1.5 second threshold). Across birds, direct responses were made to ≈ 44% of calls, which was significantly greater than chance (≈ 10%; *p* < 0.001; Fig S3 (*9*)), and this rate slightly but significantly increased over time (Fig. 1G; *p* < 0.001).

**Figure 1:**
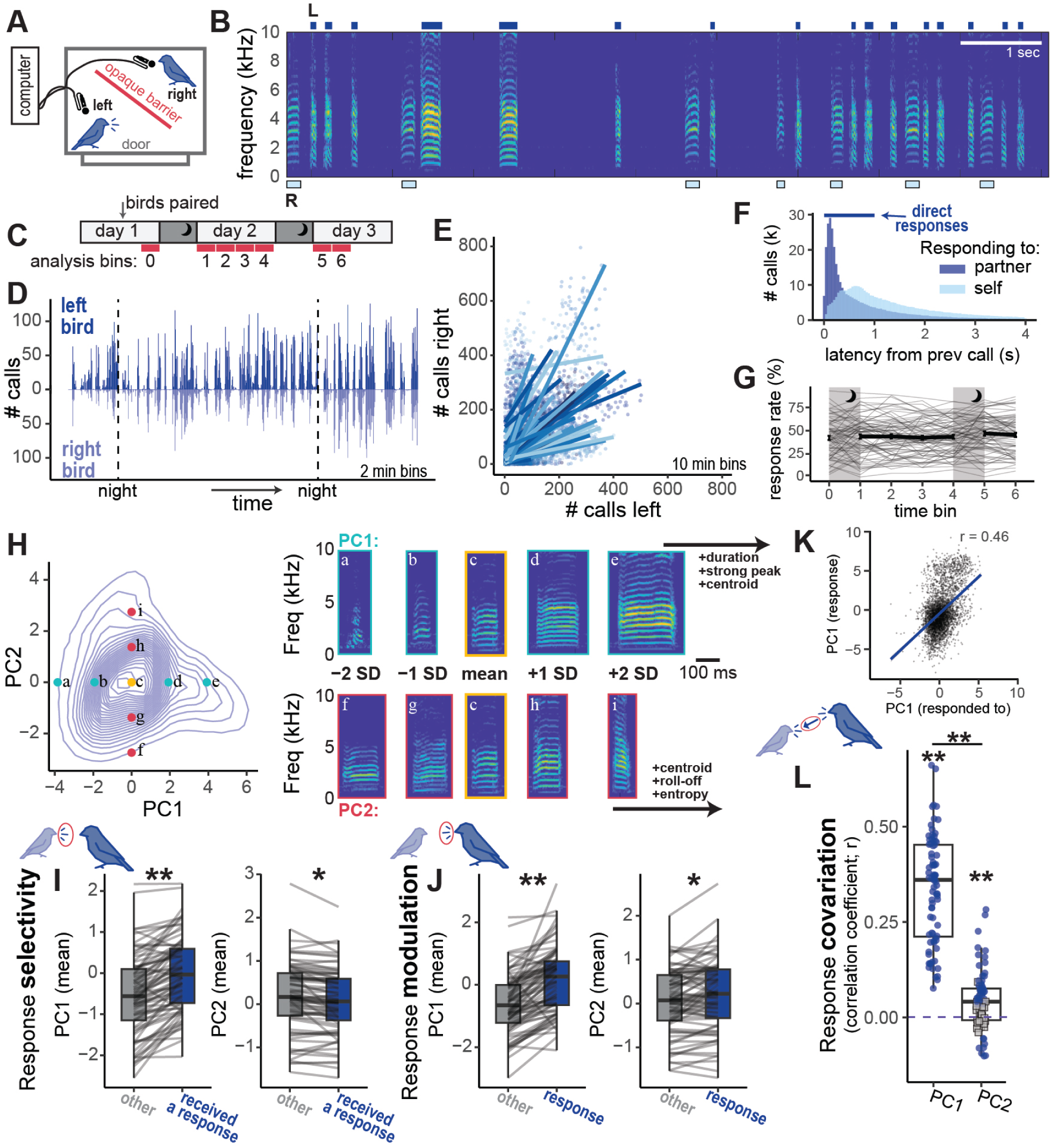
Interactions of female zebra finch pairs. A. Diagram of the recording apparatus. B. Example spectrogram of an interaction. Dark bars (top) indicate calls from the left bird, and lighter bars (bottom) indicate calls from the right bird. C. Schematic of recording timeline and bins used for analysis. D. Histograms depict the number of calls in 2 minute bins across all data analyzed for an example pair. Bars going up indicate calls from the left bird, and bars going down indicate calls from the right bird. E. Dots depict the total number of calls given by the left bird (x-axis) and right bird (y-axis) across 10 minute bins. Lines depict simple linear correlations for each pair (bins where both birds produced zero calls were not included). F. Histograms depict the response latencies for all calls that occur within 4 seconds from another call, whether a partner’s call (dark blue) or a bird’s own call (light blue). Dark blue line indicates the one second threshold to be considered a “direct response” to partner in further analysis. G. Gray lines depict the percent of a focal bird’s calls that received a direct response from the partner, and the black line depicts the mean ± sem across all birds. H. Left: density plot for all calls by PC1 and PC2, with colored dots indicating the example spectrograms to the right. Right: example spectrograms of calls along the PC1 (light blue) or PC2 (pink) axes. Text indicates the top three features loading on each of the PCs. I. Boxplots depict the acoustic features (PC1 left, PC2 right) of calls that received a response from the partner (blue) compared to other calls (gray). J. Boxplots depict the acoustic features (PC1 left, PC2 right) of calls used as direct responses to the partner (blue) compared to other calls (N). For I,J, gray lines connect the data from one bird within a recording. K. An example correlation for PC1 of direct responses to the partner (y-axis) and the calls being responded to (x-axis). L. Points depict the correlation coefficients for feature correlations between calls in response to the partner and those calls being responded to, for PC1 (left) and PC2 (right). Blue dots represent correlations that were significant within the pair (*p* < 0.05) while gray squares indicate non-significant correlations. For all panels, ∼ *p* < 0.10, ∗ *p* < 0.05, ∗∗ *p* < 0.005.

To analyze acoustic flexibility in these interactions, we measured 15 acoustic properties of all calls produced by each bird during their interactions. The first two principal components explained 25% and 13% of the variation in the data (Fig. 1H, Table 6). In general, increases in PC1 describe calls with greater “magnitude”, such as longer durations and increased relative amplitude of the dominant frequency (strong peak), whereas increases in PC2 describe brighter and less spectrally complex calls (e.g, higher frequency, higher entropy, and less filtering of upper harmonics (higher roll-off); a “brightness/spectral complexity” axis of variation); Fig. 1H). We also conducted supplementary analyses on features we were interested in *a priori* (commonly measured in birdsong; duration, entropy, dominant frequency; Fig. S1), with similar results.

We defined and analyzed three aspects of vocal flexibility. First, did individuals **selectively** respond to calls with specific acoustic features? Second, did females **modulate** features of their own calls when they produced responses? And third, on a response-by-response basis, did birds produce calls with acoustic features that **covaried** with those of her partner’s calls?

Birds exhibited selectivity in their responses to the partner’s calls on PC1 and PC2 (Fig. 1I; mixed-effects models: *p* < 0.02 for both; Table S7), driven, in part, by birds selectively responding to longer (higher PC1), lower entropy (lower PC2) calls (Fig. S1). Similarly, birds modulated the acoustic structure of their direct responses on both PCs (Fig. 1J; *p* < 0.02 for both). Finally, the acoustic structure of a response (based on both PCs) was significantly correlated with the acoustic structure of the call it was produced in response to (Fig. 1L; *p* < 0.002 for both). The correlation for PC1 was significantly stronger than that for PC2 (*p* < 0.001).

One possibility is that these changes in acoustic features of calls arise based on the use of call “types” (for example, always responding to calls of one type with that same call type). However, all three measures of vocal flexibility remained significant even when restricted to a single putative call “type” (*24, 32, 33*) (Fig. S2). Taken together, these results reveal that pairwise interactions of female zebra finches are characterized by dynamic selectivity, modulation, and covariation of call acoustic features.

### 2. Passive playbacks: birds demonstrate less vocal flexibility in response to non-contingent play-backs

To determine the degree to which vocal flexibility might characterize live interactions, we next recorded birds in two passive playback conditions. For these conditions, calls were chosen at random from an unfamiliar bird’s repertoire and played back either at (a) fixed intervals (5, 7.5, or 10 s; fixed-interval passive playback; n = 11 birds) or (b) random intervals (random-interval passive playback; n = 12 birds). Consistent with (*29*), birds readily interacted with the passive playbacks, producing more calls during playback blocks than during silence (mixed-effects models: *p* < 0.001 for both, Fig. 2A-C, Table S8). However, birds responded more slowly and less frequently to play-backs compared to a live partner. Specifically, response latencies were slower to the fixed-interval passive playback than to a live bird (Tukey’s test: *p* = 0.015) and the response rates to passive playbacks never reached the level of response to a live bird (*p* < 0.001; Fig. 2D).

**Figure 2:**
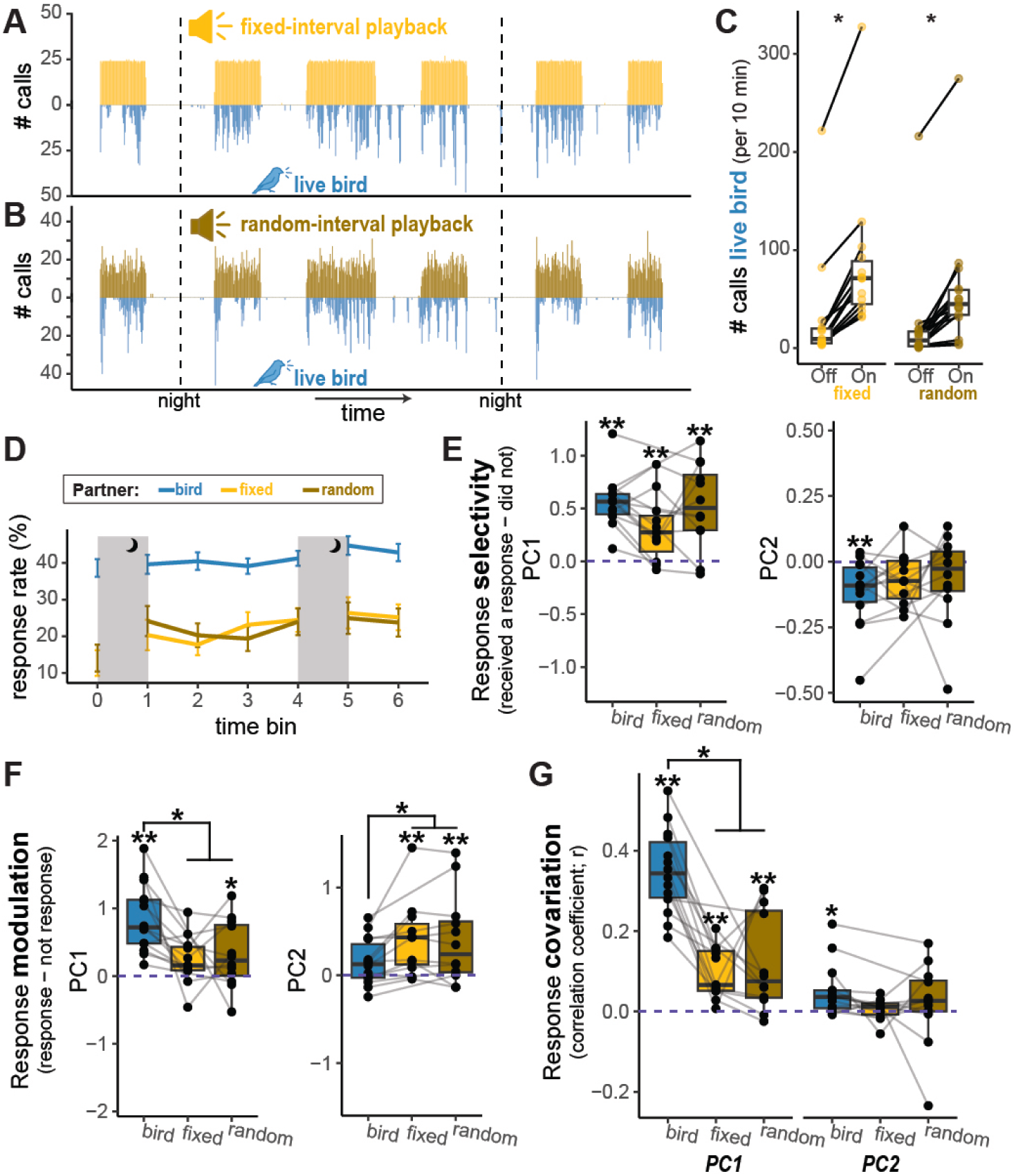
Responses of birds to passive playbacks. A,B: Histograms depict the number of calls from the passive playback (going up) and the number of calls from the bird (going down) for an example fixed-interval playback (top) and an example random-interval playback (bottom). Each bar depicts data for a 2 min bin and dashed vertical lines depict overnights (when birds to not call). The playbacks were turned on for 2 or 3 hour sessions. C) Each dot depicts the average number of calls (10 min bins) produced by a single bird during periods when the playback was off (left) and on (right), with lines connecting data from the same bird. D. Response rates across time bins. Lines depict the mean and standard error across birds. E. Response selectivity based on call features (PC1 left, PC2 right), calculated as the difference between the mean value for calls that receive a response and calls that do not receive a response. F. Response modulation based on call features (PC1 left, PC2 right), calculated as the difference between the mean value for calls that are produced as direct responses to the partner and calls that are not. G. Response covariation for PC1 (left) and PC2 (right), calculated as the correlation coefficient between all calls in response to the partner and those calls receiving a response. For E-G, each dot depicts the mean value for a bird in each condition, with lines connecting data from the same bird. For all panels, ∼ *p* < 0.10, ∗ *p* < 0.05, ∗∗ *p* < 0.005.

To investigate selectivity, modulation, and covariation to passive playbacks (acoustic flexibility), we applied the loadings from the PCA conducted during live interactions (see above) to the calls produced by the birds and passive playbacks. Birds exhibited response selectivity during live interactions and passive playbacks, in all cases responding to calls higher on PC1 (mixed effects models: *p* < 0.001 for all; 2E, Table S8). While there was no significant difference in selectivity for PC2 across conditions, selectivity on PC2 was significantly different from chance only during the live condition (*p* = 0.004; consistent with the full dataset above).

In contrast to selectivity, acoustic modulation of responses varied across conditions (mixed-effects models: *p* < 0.004 for both PCs; Fig. 2F, Table S8). Specifically, modulation of PC1 was larger during live interactions compared to each type of passive playback (*p* < 0.02 for each) while modulation of PC2 was weaker in live interactions than during playbacks (*p* < 0.004 for both). Modulation of PC1 was significantly different from chance in both the live (*p* < 0.001) and random-interval passive playback (*p* = 0.024) conditions, and modulation of PC2 was significantly different from chance only in the playback conditions (*p* < 0.003 for each).

Finally, while response covariation of PC1 was significantly different from chance when birds interacted with passive playbacks or live birds (linear models: *p* < 0.003 for all; 2G, Table S8), the magnitude of covariation was larger with a live bird than either playback (*p* < 0.001 for both). Covariation on PC2 was only significantly different from chance in the live condition (*p* = 0.013), and not during interactions with passive playbacks. Overall, we find that many aspects of the birds’ behavior differ significantly when interacting with a passive playback compared to when birds interact with a live partner.

### 3. Building and testing an interactive model (ZF-AIM) that simulates natural interactions

The previous analyses indicate that birds respond less frequently, with less modulation, and less covariation in response to passive playbacks than to a live bird. Because contingencies in responses are important in interactions, we developed ZF-AIM (Acoustic Interaction Model), an audio-LLM, to interact in real-time with a zebra finch partner. ZF-AIM used a recurrent memory transformer (*34*) to predict future states of the interaction (call timing and acoustic structure), and a neural audio codec (*35*) to generate synthetic zebra finch vocalizations based on a given bird ID. In brief, based on a sequence of vocalizations detected in an audio stream, the model predicts the time elapsed until it should produce its next vocalization. It also predicts the acoustic features of this vocalization, which are encoded in an integer call token. When the time comes to produce a vocalization, the neural codec converts this call token into an audio signal which is played from a speaker. For model training, we used audio from 34 of the 40 zebra finch pairs recorded. The remaining 6 pairs were reserved for model evaluation.

We evaluated the effectiveness of ZF-AIM using task-specific metrics, as well as through *in silico* interactions between two copies of ZF-AIM. The model was able to detect calls and assign caller identity in streaming audio with precision of 0.953 (the ratio of the number of correct detections to total predicted detections), as well as recall of 0.936 (the ratio of the number of correct detections to the ground truth number of calls). The model predicted the time until the next call more precisely (RMSE=976.0 seconds) than choosing a time interval at random from the distribution of wait times (RMSE=1126.4 seconds), predicted the identity of the next caller more accurately (Acc.=0.632) than chance (Acc.=0.496), and predicted the next call token more accurately (Acc.=0.056) than chance (Acc.=0.010). Call reconstruction was accomplished with a custom Encodec (*35*) model trained on zebra finch data. Its performance, measured with Frechet audio distance (FAD) (*36*), was lower (better) than that of neural codecs trained on general audio and on general bird audio (FAD=0.375 versus 3.306 and 7.885, respectively; see Methods).

Before recording this model with a live bird, we digitally tested it by reproducing the 40 pairs of live interactions *in silico*, in which two copies of ZF-AIM engaged in an interaction using the same bird IDs. We found that many properties of the bird-bird interaction were replicated (Fig. S4). For example, call activity was correlated across bins (*p* < 0.001) and consisted of rapid responses to the partner, with response rates of ≈ 31%. The ZF-AIMs selectively responded more to calls that were higher on PC1, and modulated their responses based on PC1 (*p* < 0.001 for both). Similar to live birds, the ZF-AIMs also covaried their responses to their partner’s calls for both PC1 and PC2 (*p* < 0.001 for both), with stronger covariation for PC1 than PC2 (*p* < 0.001).

### 4. Interactive playbacks: birds and ZF-AIM flexibly modify calling during real-time interactions

We recorded the interactions between ZF-AIM and 11 birds (Fig. 3A) and assessed how naturalistic these interactions were, first by analyzing and comparing the behavior of the birds and ZF-AIMs across interactions. As in the interactions between two birds, ZF-AIM and the bird produced correlated bursts of activity (Fig. 3B) with call numbers correlated across bins (mixed-effects models: *p* < 0.001; Table S9) and no systematic bias for the bird or ZF-AIM to produce more calls than the other (Fig. 3D). We found increased calling over time for both the bird and ZF-AIM (*p* < 0.001), and birds increased their response rate to ZF-AIM over time (*p* < 0.001), demonstrating larger changes in response rate over time than ZF-AIM (interaction: *p* < 0.001). The bird and ZF-AIM generally produced rapid responses to each other (Fig. 3E); however, ZF-AIM’s response latency was slower than the birds’ (≈ 175 ms slower; *p* = 0.018; Fig. 3G, Table S9).

**Figure 3:**
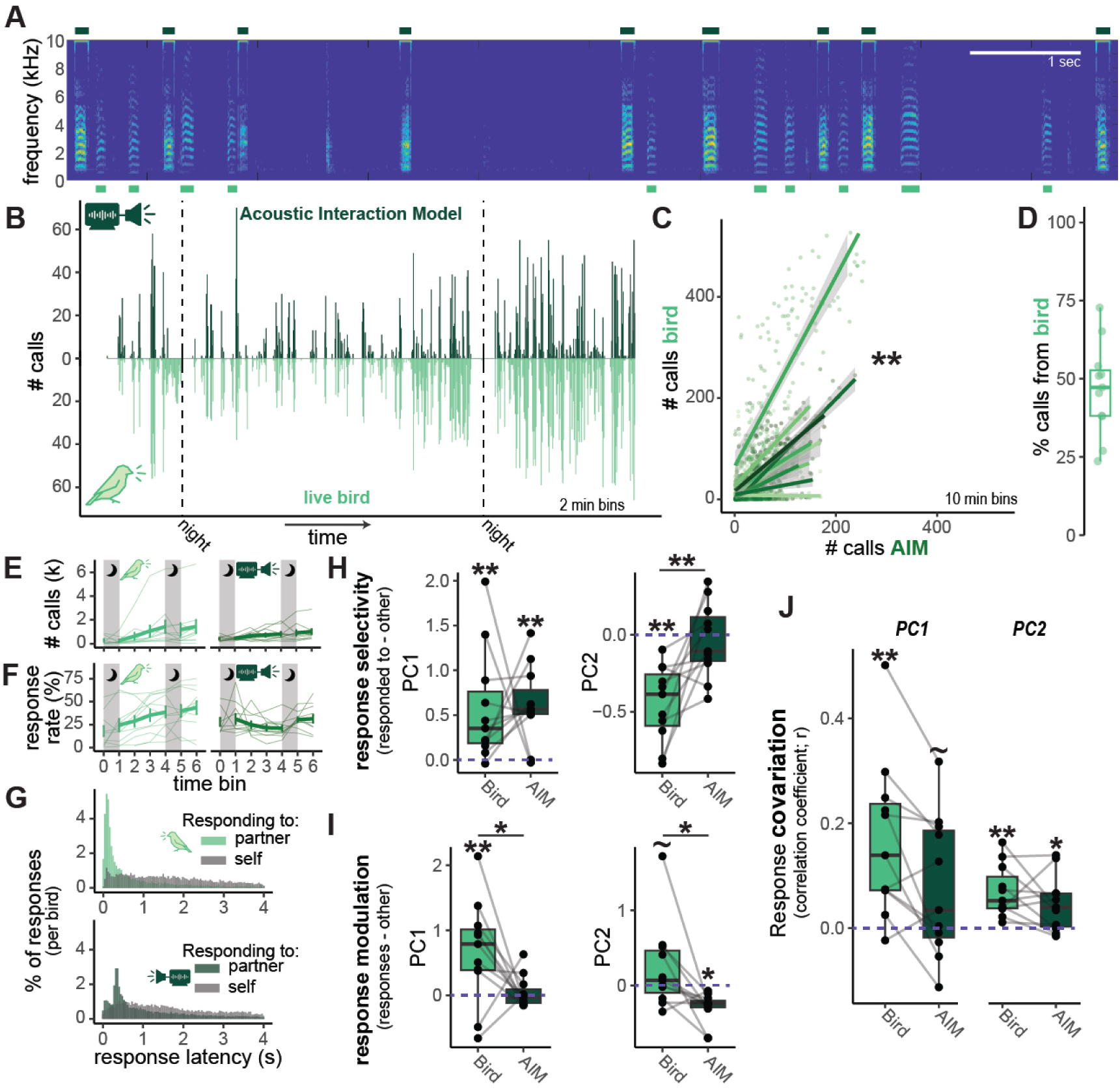
Behavior of ZF-AIM playback and zebra finches in live interactions. A. Example spectrogram showing calls from ZF-AIM playback (indicated by teal bars on top) and from the live bird (indicated by green bars on bottom). B. An example histogram showing the calling behavior of the playback (going up) and the bird (going down) across 2 minute bins. Dashed vertical lines indicate night, when birds do not call and the playback was turned off. C. Call correlations between the playback and bird. Each dot depicts the total calls produced by the model (x-axis) and bird (y-axis) within a 10 minute bin. Lines depict simple linear correlations within each pair. D. Percent of the total calls produced by the bird in each pair. E. Total calls produced across time bins by the bird (left) and playback (right). F. Response rates over time bins by the bird (left) and playback (right). For E,F, each thin line depicts data from a single bird or playback, with thick lines depicting the mean and standard error. E. Histograms depict latency of each call relative to the previous call, colored by the previous callers identity (green/teal = partner, gray = self) for birds (top) and the playback (bottom). Histograms are normalized within each bird or playback. H. Response selectivity based on call features (PC1 left, PC2 right), calculated as the difference between the mean value for calls that receive a response and calls that do not receive a response. I. Response modulation based on call features (PC1 left, PC2 right), calculated as the difference between the mean value for calls that are produced as direct responses to the partner and calls that are not. J. Response covariation for PC1 (left) and PC2 (right), calculated as the correlation coefficient between all calls in response to the partner and those calls receiving a response. For H-J, each dot depicts the mean value for a bird or a playback, with lines connecting data between a bird and the playback she received. For all panels, ∼ *p* < 0.10, ∗ *p* < 0.05, ∗∗ *p* < 0.005.

We found patterns of selectivity, modulation, and covariation, both by the birds and ZF-AIM, despite the differences in response timing. Both ZF-AIM and birds selectively responded to calls higher in PC1 (mixed-effects models: *p* < 0.002 for both), while only the birds selectively responded to calls lower in PC2 (birds: *p* < 0.001, ZF-AIM: *p* = 0.596; Fig. 3H). Consistent with these patterns, we found no difference between ZF-AIM and bird for PC1 selectivity (*p* = 0.850), but did find a difference for PC2 selectivity (*p* < 0.001). Birds and ZF-AIM differed in their modulation of PC1 and PC2 (PC1: *p* = 0.021; PC2: *p* = 0.009; Fig. 3I, Table S9), driven by birds producing responses with higher PC1 and marginally higher PC2 values (*p* = 0.001 and 0.084, respectively) while ZF-AIM produced responses lower on PC2 (*p* = 0.033). Finally, birds and ZF-AIM exhibited similar response covariation, with no significant difference in covariation for either PC (*p* > 0.15 for each; Fig. 3J, Table S9). Birds demonstrated significant covariation for PC1 and PC2 (*p* < 0.001 for both), while ZF-AIM demonstrated significant covariation for PC2 (*p* = 0.007) and marginally non-significant covariation for PC1 (*p* = 0.085). Overall, despite some differences in the timing and acoustic modulation of calls produced by the birds and ZF-AIM, we observed bursts of acoustic interaction with dynamic flexibility in acoustic features from birds and ZF-AIM.

### 5. Model comparison: Reduced flexibility from the model impacts only some aspects of a bird’s vocal flexibility

We conducted an additional playback experiment to determine the importance of acoustic flexibility exhibited by ZF-AIM for naturalistic behaviors from the birds. Specifically, ZF-AIM-ablated differed from ZF-AIM in that it ignored the acoustic properties of the live partner’s calls, and selected its own call token at random (see Methods; n = 11 birds). Therefore, the ablated model retained ZF-AIM’s call timing and response behaviors, while drastically reducing the ability of ZF-AIM-ablated to exhibit any measure of flexibility based on acoustic properties. Indeed, ZF-AIM-ablated and birds produced correlated bouts of calling (*p* < 0.001), with drastically reduced selectivity, modulation, and covariation from the model (see Fig. S5).

Similar to the interactions with passive playbacks, zebra finches exhibited lower response rates to both ZF-AIM and ZF-AIM-ablated relative to a live bird at the beginning of the interaction. However, unlike passive playback interactions, the response rates to both ZF-AIM and ZF-AIM-ablated were similar to the response rates to a live bird by the end of the second day (Fig. 4A). Response latencies were similar in all three conditions (mixed-effects model: *p* = 0.654).

**Figure 4:**
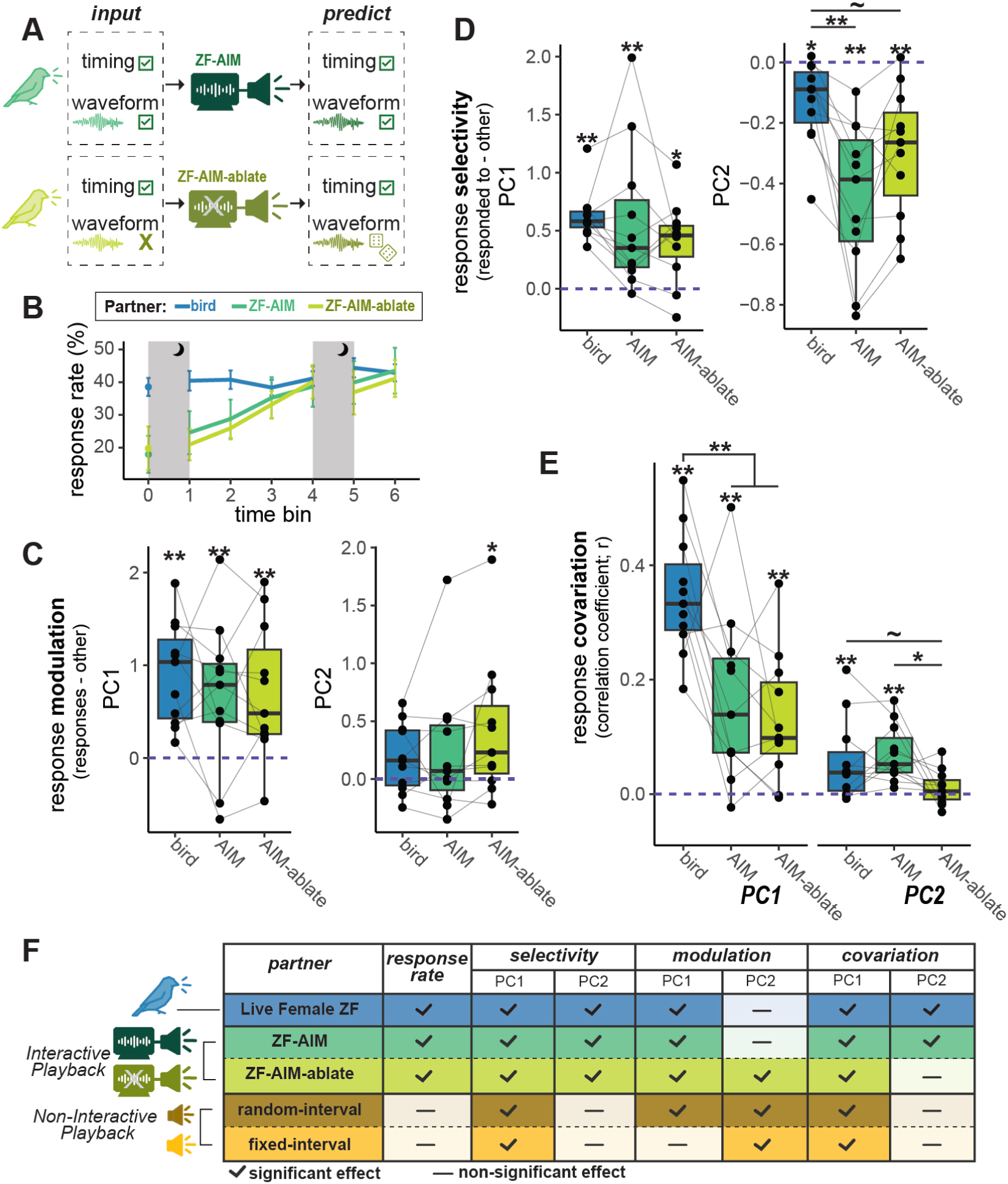
Responses of birds to interactive playbacks. A. Diagram explaining our two interactive models. B. Response rates to the partner across time bins. Lines depict the mean and standard error across birds in each condition. C. Response selectivity based on call features (PC1 left, PC2 right), calculated as the difference between the mean value for calls that receive a response from a live bird and calls that do not receive a response from a live bird. D. Response modulation based on call features (PC1 left, PC2 right), calculated as the difference between the mean value for calls that are produced as direct responses to the partner and calls that are not. E. Response covariation for PC1 (left) and PC2 (right), calculated as the correlation coefficient between all calls in response to the partner and those calls receiving a response. F. Table depicting the pattern of significant effects for each condition. Analyses are limited to birds that were used in playback experiments (this Fig. and Fig. 2). Note that the check marks for response rates refer to whether response rates were non-significantly different from those during live interactions by the end of the interaction. For C-E, each dot depicts the mean value for a live bird, with lines connecting data from the same bird in different conditions. ∼ *p* < 0.10, ∗ *p* < 0.05, ∗∗ *p* < 0.005.

Birds selectively responded to calls from their partner, regardless of whether the partner was a live bird or an interactive model (ZF-AIM or ZF-AIM-ablated) based on both PCs (linear models: *p* < 0.03 for all; Fig. 4B, Table S10). Selectivity based on PC1 features was similar with live birds and the interactive models. However, for PC2, the magnitude of selectivity was significantly larger during interactions with ZF-AIM than a live bird (*p* = 0.001), and marginally larger during interactions with ZF-AIM-ablated than a live bird (*p* = 0.082).

Birds modulated their responses along PC1 regardless of the partner (linear model: *p* < 0.005 for all; Fig. 4C, Table S10), similar to what we observed during passive playbacks. When birds interacted with ZF-AIM-ablated (but not a live bird or ZF-AIM), they modulated their responses for PC2 (*p* = 0.015), with a trend toward this modulation being larger in magnitude than during live interactions (mixed-effects model: *p* = 0.092).

Finally, we found that the magnitude of feature covariation in responses varied by interaction partner (Fig. 4D, Table S10). While birds exhibited covariation of PC1 in all three conditions (linear models: *p* < 0.002), covariation was stronger with a live partner than with either AIM (*p* < 0.003 for each). Covariation of PC2 was significant only with a live partner or ZF-AIM (*p* < 0.002 for each), but not with ZF-AIM-ablated (*p* = 0.522). The magnitude of covariation was significantly lower during interactions with ZF-AIM-ablated than with ZF-AIM (*p* = 0.030) and marginally lower than with a live bird (*p* = 0.096).

## Discussion

By integrating large-scale analysis and novel AI-based interactive playbacks, we discovered principles underlying vocal interactions and factors that regulate natural discourse structure. During dyadic interactions, female zebra finches exhibited robust, rapid, selective, and modulated responses to the vocalizations of their partner, and these responses covaried with the acoustic properties of their partner’s calls. These patterns resemble vocal flexibility observed in human conversations, such as intonation and style matching (*5–7*). Intriguingly, this flexibility in female finches exists even though the acoustic structure of female calls is not learned (*37, 38*), suggesting that similar modulations may occur in other species, regardless of their vocal learning abilities.

We used passive and interactive playbacks to test the features that elicit naturalistic vocal flexibility from a live bird. At a basic level, our passive playbacks examined the importance of contingency (e.g., call timing and structure conditioned on interaction history) by testing how birds responded to a natural repertoire of calls but in which the timing and acoustic structure were independent of the bird’s behavior. In contrast, our generative language model (ZF-AIM: Acoustic Interaction Model) used its own prior behavior and the behavior of a live bird to determine the timing and acoustic structure of the next call in the interaction. Finally, we developed ZF-AIM-ablated as an intermediate test, where, like ZF-AIM, the timing of calls was predicted based on the interaction, but, like the passive models, the call selection was made without knowledge of any call acoustics (from either the bird’s calls or the model’s calls).

These comparisons revealed principles underlying natural interactions. First, interactive contingency of call timing was key for increasing and maintaining responses. Birds interacting with passive models never achieved response rates equivalent to those seen in natural interactions, while birds increased response rates to interactive models over time. Second, regardless of the partner, birds selectively responded to longer/stronger calls (high on PC1) and often produced similarly longer/stronger calls in response. Zebra finches are a highly social species, and this acoustic variation likely relates to call function (*24, 32, 33, 39*). Finally, acoustic flexibility from the partner appears important for subtle modulations of the brightness/spectral complexity (PC2) of calls. Birds covaried but did not modulate this aspect of their call when interacting with a live bird or ZF-AIM. In contrast, birds modulated (increased) but did not covary the brightness/spectral complexity features of their calls when responding to passive playbacks and ZF-AIM-ablated. Indeed, despite some variation in the magnitude of effects, ZF-AIM was the only playback to achieve the same pattern of significant effects as a live interaction (Fig. 4F).

It is notable that, while ZF-AIM did not always correctly predict the next call when tested on interactions between two birds, the conversations it produced with live bird partners were largely naturalistic. This stands in contrast to the behavior of some early chatbots, where errors in predicting the next word could lead to conversational breakdowns (*40*). One especially intriguing possibility is that the external feedback from a partner (whether a live bird or *in silico* ZF-AIM) was sufficient to correct the interaction when the model did not act like a bird would. Alternatively, it is possible that the interactions appear naturalistic because errors made by the model were small or because our measurements miss some non-naturalistic aspect of the interaction with ZF-AIM. Further experiments could help explain model performance and elucidate how robust zebra finch interactions are to aberrant responses by one interlocutor. Additionally, ablation studies can test how these measures vary in relation to other computational aspects such as the context length (which can influence predictive capability; e.g., sperm whales (*41*), marmosets (*42*), and Bengalese finches (*43*)).

Taken together, by coupling analyses of naturalistic interactions with AI-driven experimental manipulation, we provide a scalable and generalizable approach for discovering fundamental patterns of acoustic interactions across species and contexts. Our comparison of an AI-based interaction model to passive playbacks and ablated models revealed the importance of temporal contingency and acoustic flexibility on naturalistic interactions. Assembling an analysis framework to quantify the “naturalness” of interactions is challenging (*44*), and we propose that our framework incorporating measures of response rates, timing, selectivity, modulation, and covariation comprehensively characterizes natural vocal interactions and can be applied to a wide range of species and scaled up to the analysis of larger group dynamics (*4, 22, 24, 27, 39*).

## Acknowledgements

We thank the members of the Woolley and Sakata labs for help with bird care and feedback on analysis. We thank the staff of the Earth Species Project for help with logistical support, fundraising, and feedback on the project. We thank Felix Effenberger, David Robinson, Brittany Solano, and Maddie Cusimano for valuable feedback on the manuscript.

## Author contributions statement

Conceptualization: LSJ, BH, JL, MM, AR, JTS, EC, OP, SCW. Data curation: LSJ, BH, JL, MM, MA. Formal analysis: LSJ, BH, JL, MM. Funding acquisition: AR, JTS, OP, SCW. Investigation: LSJ. Methodology: LSJ, BH, JL, MM, JTS, EC, OP, SCW. Resources: LSJ, MA, JTS, EC, OP, SCW. Software: LSJ, BH, JL, MM, MA. Supervision: MG, JTS, EC, OP, SCW. Validation: LSJ, BH, JL, MM, EF. Visualization: LSJ, JL, DK. Writing (original draft): LSJ, BH, JL, MM, JTS, EC, SCW. Writing (review and editing): LSJ, BH, JL, MM, EF, DK, JTS, EC, OP, SCW.

## Data availability

All data required to reproduce the statistical analyses in the paper (i.e., extracted timestamps and acoustic features of all calls) will be available upon acceptance. However, as in the human realm (*45*), introducing LLM’s to animals raises new ethical considerations (*46–48*). While we carefully limited potential harms in this study (e.g., using captive animals whose basic needs were met and using relatively brief experimental exposures), the involvement of a black-box AI increases the risk of inadvertently playing sounds that may negatively impact individuals or populations, particularly in the wild (*46*). At the same time, we highlight that sophisticated modeling and experiments with “digital twins” may reduce the burden placed on laboratory animals and allow additional insights into species not amenable to experimental conditions (*49*). In light of these considerations, we do not release model code or weights at this time.

## 1 Materials and Methods

### 1.1 Data Collection

#### 1.1.1 Animals

Female zebra finches were raised in a breeding colony at McGill University with parents and siblings until 60 days of age and then housed in same-sex group cages within the colony. All birds were housed on approximately 14L:10D or 15L:9D light cycles with water and food (seed) provided *ad libitum*. All birds in the experiment were adults (> 4 months) and housed in same-sex group cages in sound attenuating chambers (TRA Acoustics, Ontario, Canada).

Animal care and procedures followed all Canadian Council on Animal Care guidelines and were approved by the Animal Care Committee of McGill University.

#### 1.1.2 Recordings

For recordings, birds were each placed into their own individual cage (8” cube) within a sound attenuating chamber (26” x 21” x 19”; TRA Acoustics). Visual contact was obscured using an opaque barrier because visual contact is known to influence calling behavior (*31*). Birds were paired in the afternoon and recorded for at least 48 hours. Birds in all pairs had been isolated from each other for at least six months.

Live interactions were recorded using two omnidirectional microphones (Countryman Associates, Inc), each positioned above a bird’s cage, and digitized at 44.1 kHz. Initial recordings (Pairs 1-26) were collected using Sound Analysis Pro (*50*) with each microphone recording to a separate mono track in parallel. However, we discovered that this method sometimes resulted in slight mis-alignment between the two channels (see below). Subsequently, all recordings were collected using Audacity ^1^ (versions 3.3.3, 3.4.2, or 3.5.1) in stereo with the microphone above each bird recording to separate channels simultaneously.

To record birds interacting with playbacks, we replaced the left cage with a set of speakers (Logitech), but kept the same visual barrier in place. The left microphone remained positioned slightly in front of the speakers. For playbacks, we calibrated the speakers to the peak amplitude of the playback call with the median amplitude across all calls in that playback’s call repertoire (see below). Specifically, the call with the median amplitude was set to a peak relative amplitude of 0.1, and all calls were normalized accordingly. We then generated a calibration tone with a relative amplitude of 0.1, and set the speakers to an amplitude of approximately 83.9 dB (± 1.2 SD) at the center of the live bird’s cage.

### 1.2 Data Processing

#### 1.2.1 Initial processing

The initial 26 recordings made in Sound Analysis Pro consisted of separate 33-second mono recordings for the left and right channels. In some cases audio samples were dropped from one recording and not the other, causing some pairs of left and right recordings to be misaligned in time. Based on the recording start time metadata and recording durations, we identified pairs of left and right recordings which had a misalignment of at least 50 milliseconds. These 33-second recording pairs were excluded from further analysis (27706 recording pairs excluded from a total of 136145 recordings, median 1215.5 recording pairs excluded per 48-hour recording session). Each of the remaining, non-excluded recording pairs, were combined into one 33-second stereo audio recording for subsequent analyses. For computing latencies between vocalizations, we accounted for the missing data: if a vocalization *x* occurred, and before the next recorded vocalization there was an excluded recording, then the time until the next vocalization was marked as unknown and *x* was omitted from analyses relying on inter-vocalization latencies.

Subsequent zebra finch pairs, and all playback experiments, were recorded in Audacity. Each of these resulted in a single multi-day stereo recording, and did not suffer from misalignment. These were split into 30-second clips for subsequent analyses. To avoid the confound of missing data, we limited our analysis of changes in calling over time to pairs without missing data (27–40). We note that limiting our analyses to pairs 27-40 does not change any of our primary results.

#### 1.2.2 Detection and diarization of vocalizations

As an initial data processing step, it was necessary to detect the vocalizations recorded by the stereo microphones and identify the bird that produced each vocalization. To do so, we developed a machine learning model using the open-source software Voxaboxen (*51*), which we call ZFVoxaboxen. ZFVoxaboxen predicts bounding boxes (i.e. start- and stop-times) of vocalizations in the stereo audio. For each bounding box, ZFVoxaboxen also predicts which bird (Left or Right) produced the boxed vocalization.

To develop ZFVoxaboxen, two of us (BH and LJ) annotated twelve 33-second recordings, from each of 13 recorded pairs. Annotations were done in Raven^2^ and included start and end times of each vocalization, as well as the identity of the vocalizer (Left or Right). Identity determinations were made based on annotators’ perception of the source in the stereo recordings and visualization of the relative amplitude in each microphone using spectrograms. In general it was very easy to tell which bird was vocalizing, based on relative amplitude of the calls in the two audio channels. Overall, we annotated 4275 vocalizations across all 156 recordings. On average, annotated vocalizations were 0.136 seconds long (standard deviation = 0.042). We split these annotated data into a train set, consisting of eight bird pairs (Pairs 1-5, 11-13), and a test set, consisting of the remaining five pairs (Pairs 6-10). To double the size of our train set, we created a channel-swapped version of each recording.

ZFVoxaboxen is a deep neural network. It consists of a feature extractor, followed by a final prediction layer. For the feature extractor, we use AVES-bio (*52*), which is a transformer-based architecture that was pre-trained on 360 hours of animal audio. Although AVES-bio was pre-trained on audio at 16 kHz, we found in initial experiments that providing ZFVoxaboxen with audio at 32 kHz (effectively playing at half speed) gave better results. After being re-sampled to 32 kHz, each channel of the stereo audio is passed in to the feature extractor, which produces feature vectors (each 768-dim) at a frame rate of 100 Hz. At each frame, we concatenate the feature vectors coming from the two stereo channels, producing a 1536-dimensional feature vector. This feature vector is passed into a final prediction layer as described in Mahon et al. (*51*). We do not use Voxaboxen’s bidirectional prediction option described in section 3.2 of (*51*), since it was developed after we created ZFVoxaboxen.

We trained ZFVoxaboxen as in (*51*) for 25 epochs. During training, we divided the recordings into short windows with 50% overlap between windows. Based on initial experiments, we found that training with a low batch size improved results; therefore, ZFVoxaboxen was trained using a batch size of 1. The parameters of the AVES feature extractor were unfrozen after the first three epochs. We performed a grid search across the learning rate and window duration hyperparameters, using the procedure described below. Learning rate was chosen from {.0001, .00006, .00003}, and window length was chosen from {2, 4} seconds. Training took approximately two hours on one Nvidia A100 GPU.

Bounding boxes are constructed from model outputs as in (*51*). To evaluate the performance of ZFVoxaboxen, we follow a standard sound event detection evaluation procedure (*51*). First, we match each predicted bounding box to at most one annotated bounding box, using an intersection-over-union (IoU) threshold of 0.5. Once box predictions are matched with annotations, for both Left and Right birds, we compute classification precision, recall, and F1 scores. We then compute class-averaged (i.e. macro-averaged) precision, recall, and F1 scores, by taking the average of these scores across both categories (Left and Right).

To tune the learning rate and window length hyperparameters, we did the following. From the 8 pairs in the train set, we set aside four recordings (plus their channel-swapped versions) as a validation set, which we used to compare different hyperparameter settings. We trained versions of ZFVoxaboxen using the remaining recordings, and selected the hyperparameters that yielded the best macro F1 performance on the validation set. We trained the final version of ZFVoxaboxen using the selected hyperparameters on the entire train set. On the test set, ZFVoxaboxen achieved a macro F1 score of .956 at 0.5 IoU, which is very close to the optimal F1 score of 1.0. It had a macro precision of .958 and a macro recall of .955.

#### 1.2.3 Vocalization denoising

We created a denoised version of each recording, in which the audio signal corresponding to the zebra finch vocalizations was isolated and the noises coming from other sources (e.g. fan noise, wing flaps) were discarded. These denoised vocalizations were used in subsequent steps: as a call bank (for passive playbacks), as training data for the ZF-AIM-encoder and ZF-AIM-decoder (described below), and for computing acoustic features (e.g., dominant frequency and spectral entropy).

We rely on 4-source BirdMixit (*53*) for denoising. BirdMixit is a model trained to separate mono recordings of birds into four audio tracks (called *stems*), each corresponding to a different audio source (multiple stems may just contain noise). To denoise a stereo recording, we denoised each channel separately. For each channel, we first silenced all audio outside of the detected vocalizations and then we passed the audio into the BirdMixit model. For each stem, we used ZFVoxaboxen to detect any zebra finch vocalizations in that stem, and silenced all sounds which were not detected as zebra finch vocalizations. Finally, the four stems were re-combined into one mono recording, which was the final denoised audio for the channel. To evaluate denoising quality, we inspected a selection of 60 randomly chosen recordings from 5 pairs, each containing at least one vocalization (575 total calls). We found no instances of acoustic artifacts from the denoising.

#### 1.2.4 Measurement of acoustic features

We measured 15 acoustic features calculated using the Python library Essentia (*54*). These features were: 1. Duration (s) 2. Dominant frequency (Hz) 3. Shannon entropy (call ‘noisiness’ relative to a pure tone) 4. Dominant frequency slope (Hz) 5. Dominant frequency standard deviation (Hz) 6. Inharmonicity (from spectral peaks) 7. Attack time (log of the attack time of the signal envelope) 8. Strong peak (relative strength of the dominant frequency) 9. Flatness (ratio between the geometric mean and the arithmetic mean of the amplitude envelope) 10. Flatness time (change in flatness over time) 11. Roll-off (frequency cutoff maintaining 85% of the power) 12. Spectral centroid (Hz) 13. Spectral complexity 14. Derivative (time derivative of the amplitude envelope) 15. Dissonance (from spectral peaks). Prior to measuring acoustic features, we low pass filtered at 8 kHz.

#### 1.2.5 Call classification

We clustered calls into potential call “types” to test whether the acoustic modulation observed may be primarily due to switching among distinct call types. Specifically, we aimed to limit our acoustic analyses to a single call type to see whether we continued to observe acoustic flexibility. To this end, we utilized a dataset of adult zebra finch vocalizations with call types assigned (*24, 32, 33*). We trained a logistic regression classifier based on the following acoustic features: flatness, spectral centroid, entropy, roll-off, strong peak, complexity, dominant frequency, mean dominant frequency, dissonance, and tristimulus. To better model the female zebra finch interaction we restricted the call types referenced in (*32*) to DC, Di, LC, Te, Th, Wh. These classes are associated with contact, distress, and alarm calls. Our final goal is not to predict the context or behavior but to have a way of looking at acoustic properties of well-established call types. We used the trained logistic regression classifier to infer call types for all calls processed in our dataset.

### 1.3 Passive playbacks

For passive (i.e. non-interactive) playbacks, we played back randomly chosen calls at fixed intervals (fixed-interval passive playback) or at randomly-chosen intervals (random-interval passive playback). For fixed-interval passive playback, the interval was fixed at 5, 7.5, or 10 seconds for a given bird (roughly matching the average rates of call production in live birds). For random-interval passive playback, the interval between two successive calls was drawn randomly with probability to reproduce the same total call production as one of the metronome intervals (i.e., 5, 7.5 or 10 seconds between calls, on average); one bird’s playback was set at an average of 6.67 seconds. For both passive playback settings, calls were played for 2 or 3 hour blocks, separated by at least 2 hours of silence (3 blocks per day). For each playback recording, a bird ID was chosen and the calls produced by the playback were randomly chosen from a call bank associated with that bird ID. The call bank consisted of 1000 randomly chosen calls from each bird ID, denoised using the procedure described above. The bird ID selected for the playback was always unfamiliar to the focal bird, emulating the experimental design for interactions between live pairs of birds.

### 1.4 ZF-AIM

We propose a modeling pipeline that aims to replace one bird in a vocal interaction between two birds. Specifically, we develop an audio-LLM that detects calls made in an interaction between two zebra finches, predicts the next calls that will be made, and generates the audio of these calls. We designed ZF-AIM to be used in a live interaction setting where audio is streamed in small audio segments. An illustration of the workflow for live interaction is shown in Fig. 5 and a detailed summary is given in Table 1.

**Figure 5:**
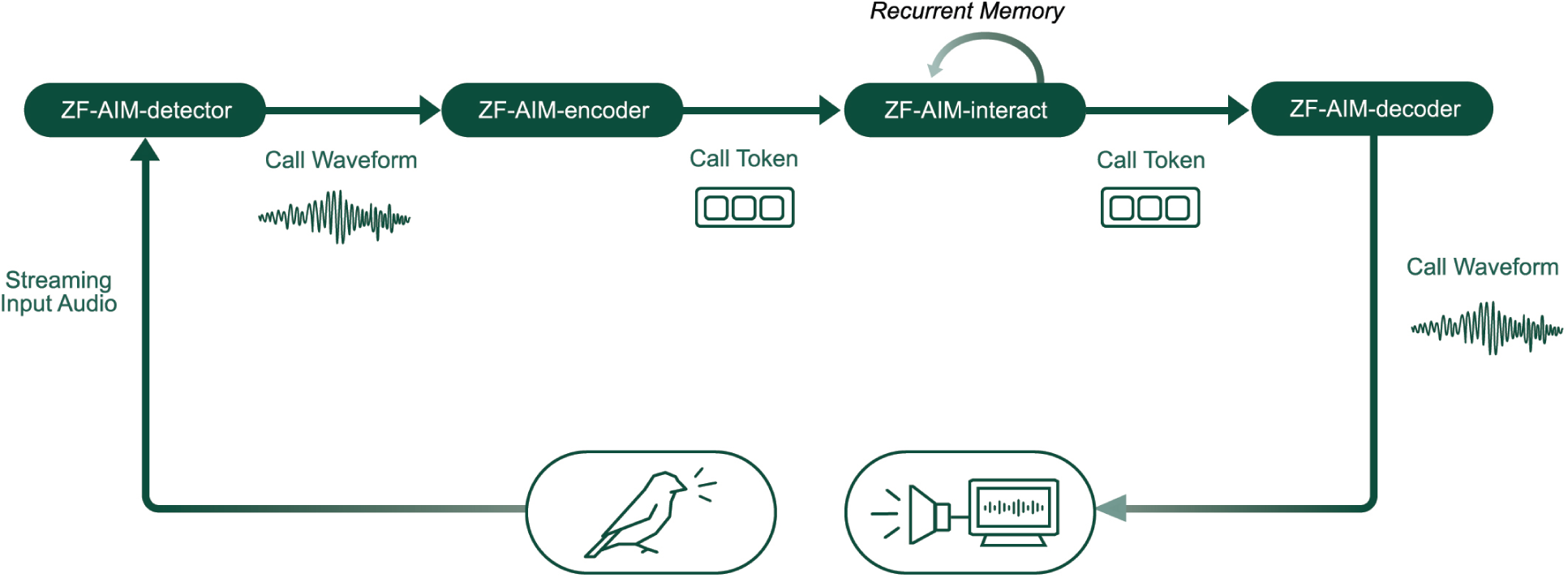
Live interaction workflow.

**Table 1:** Model inputs and outputs, in order. The sequence of events seen by the interaction model begins with a *START* event and then alternates between two types of events, *WAIT* events and *CALL* events.

| Model | Input | Output |
| --- | --- | --- |
| Live Detection Model | Streamed waveform (SR: 32k Hz)<br>Cropped call<br>Call timestamp | Frame-level detection bits (SR: 50 Hz) |
| Call Acoustic Encoder | Call waveform (SR: 16k Hz) | Sequence of acoustic feature vectors (SR: 50 Hz)<br>Sequence of multi-layer acoustic tokens (SR: 50 Hz) |
| Call Token Encoder | Sequence of acoustic feature vectors (SR: 50 Hz) | Call token |
| Interaction Model | Sequence of events<br>Generator bird ID<br>Partner bird ID<br>Daytime (True/False)<br>Days since the trial starts | Next event in sequence |
| Call Token Decoder | Call token | Sequence of multi-layer acoustic tokens (SR: 50 Hz) |
| Call Acoustic Decoder | Sequence of multi-layer acoustic tokens (SR: 50 Hz)<br>Bird ID | Call waveform (SR: 16k Hz) |

There are several sub-models involved in ZF-AIM: First, in order for ZF-AIM to engage in a real-time interaction, it required a call detection model that operates on-line as audio streams in. The model used for off-line data processing (ZFVoxaboxen) relies on non-causal attention to accurately predict call offsets, so does not adhere to these constraints. Therefore, we developed an **on-line call detection model** (ZF-AIM-detector).

Once a call is detected, the variable-length call waveform is converted to an integer call token using ZF-AIM-encoder, the **call encoder**. Specifically, the call encoder predicts a probability distribution over a learned codebook, and a call token is sampled according to this distribution. This discrete, fixed-length representation was necessary for the event-based modeling we describe in the next paragraph.

To model vocal interactions, we treat calls as discrete events, separated by waiting periods. Similar event-based modeling frameworks have been used widely in midi music generation (*55–57*). More precisely, our **interaction model** ZF-AIM-interact is fed a sequence of *WAIT* and *CALL* events. Each *WAIT* event consists of a duration of silence, and each *CALL* event consists of the caller identity and an integer *call token* which encodes the acoustic properties of the call. Then, ZF-AIM-interact predicts a probability distribution over the next wait duration, next call token, and next caller identity. A sample is drawn according to each of these distributions, which together constitute the next *WAIT* and *CALL* events.

In a live interaction with a bird, if ZF-AIM-interact predicts that the next call comes from itself, the call tokens predicted by ZF-AIM-interact are converted back to a waveform using ZF-AIM-decoder, the **call decoder**. This audio is played back to the bird through a speaker.

#### 1.4.1 Model Architecture and Data

Throughout, we use two model architectures: Transformers with recurrent memory^3^ (*34*) and Encodec models^4^ (*35*). We adopted transformers for sequence-to-sequence tasks, given their successful application to similar problems, including next syllable prediction in Bengalese finches (*43*). Since we required ZF-AIM to interact in real-time with a zebra finch, we adopted recurrent memory transformers, specifically, because they were able to model arbitrarily-long context lengths without an increase in processing time. This is in contrast to the situation with a typical transformer, which has processing time that scales quadratically with the context length. We adopted Encodec for audio compression because it exhibited a good trade-off of compression, latency, and reconstruction quality, and because it had officially supported code repository.

For recurrent memory transformers, based on observing a good trade-off of speed and performance in initial experiments, we used 3 attention layers with 16 heads, head dimension 16 for each head, and 10 memory tokens. Throughout, models were randomly initialized at the beginning of training.

We used our entire dataset of natural interactions (Pairs 1-40) as the development set and split the recordings into the training set (Pairs 1-34) and the validation set (Pairs 35-40). Since we had already held out Pairs 6-10 to evaluate ZFVoxaboxen, for the purpose of evaluating ZF-AIM-detector we additionally held out Pairs 6-10.

#### 1.4.2 ZF-AIM-detector

The live detection model ZF-AIM-detector consists of the encoder from a pre-trained Encodec model, followed by a detection head. The detection head is a recurrent memory transformer which predicts, for each frame and each caller ID (self/partner), if it is part of a call (0 or 1). To detect an entire call, we simply find the locations of changes: a call start if the detection bit changes from 0 to 1, and a call end if the detection bit changes from 1 to 0. The signal between the detected start and end is cropped as a detected call.

##### Training

The Encodec feature extractor of ZF-AIM-detector was pre-trained on the xeno-canto (*58*) portion of the pseudo-vocalization dataset from (*59*). Note that, below, we describe an Encodec model pre-trained entirely on zebra finch audio. In preliminary experiments we observed that using the Encodec pre-trained on only zebra finch audio led to degraded detection performance; we therefore adopted a larger pre-training dataset here.

For the Encodec model hyperparameters, we followed closely the setting file provided by the Audiocraft repository^5^. The only changes are that we used 8 layers of Residual Vector Quantization (RVQ) instead 32 layers, and we used [8, 5, 4, 4] downsampling ratios instead of [8, 5, 4, 2] which yielded a frame rate of 50 Hz when trained with 32kHz audio signals. The Encodec feature extractor was pre-trained for 50 epochs, using the default training settings from Audiocraft^6^.

For training ZF-AIM-detector, the audio recordings were chunked into 5-sec sub-clips with 5-sec hop, and then grouped into mini-batches of size 5. ZF-AIM-detector was trained for 50,000 steps with the Adam (*60*) optimizer and a learning rate of 1e-4. Hyperparameters were tuned in initial experiments. Binary cross entropy was used as the loss function. During training, the weights of the Encodec feature extractor were unfrozen.

When ZF-AIM-detector is used as a component of ZF-AIM in interactive playbacks, the streaming input audio segment is 0.1-sec long. To reflect this setting during training, we further segmented a 5-sec sub-clip into 0.1-sec sub-segments which were fed to the Encodec model for feature extraction. The sequence of features extracted by Encodec are then passed to the detection head.

We first trained a version of ZF-AIM-detector with the human-annotated data used to train ZFVoxaboxen. However, we found that it only achieved 0.826 macro F1 (compared to F1 of 0.956 for ZFVoxaboxen on the same dataset). To improve performance, we increased the amount of training data by using call detections produced by ZFVoxaboxen (Section 1.2.2). We used detections for all recordings from Pairs 1-5 and 11-34, a total of 487 hours of audio (containing total of 34 hours of calls). Final metrics are reported below.

In live interactions, we observed that ZF-AIM had a strong tendency to respond to itself repeatedly, likely due to incorrectly detecting its own calls as coming from the partner. To avoid this, we employed a method described in (*29,30*): a 10 kHz sine wave tone with magnitude 0.1 was added to the calls produced by ZF-AIM. Since 10 kHz is above zebra finches’ auditory capability (*61*), this added “audio fingerprint” is not perceived by the bird. Whenever ZF-AIM-detector detects a call, the first 0.05 seconds of call audio are converted to a frequency representation |*X* |^*start*^ using a fast Fourier transform followed by taking an absolute value. The magnitude 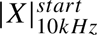 at 10kHz is compared to the maximum magnitude across all frequencies 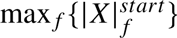. This is repeated for the last 0.05 seconds of call audio, yielding 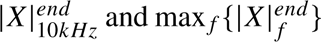. The caller ID is reassigned to be 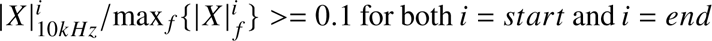, and to be the partner otherwise.

After these detection steps are performed, call audio is downsampled from 32 kHz to the 16kHz expected by ZF-AIM-encoder. This removes the 10kHz tone, which is above the Nyquist frequency of audio at 16kHz.

##### Evaluation

The evaluation process for ZF-AIM-detector is the same as described in Section 1.2.2. Without the 10 kHz tone, ZF-AIM-detector achieved, at 0.5 IoU, a 0.944 macro F1 score, 0.953 macro precision, and 0.936 macro recall, which are close to the performance of ZFVoxaboxen.

With the 10kHz tone used to re-assign caller ID predictions, ZF-AIM-detector achieved, at 0.5 IoU, a 0.934 macro F1 score, 0.948 macro precision, and 0.922 macro recall (for this, the 10 kHz tone was added post-hoc to the detection test set, which consisted of bird-bird interactions only). Qualitatively, during interactive playbacks, the version of ZF-AIM using the 10 kHz tone to assign caller ID did not suffer from the problem of making repeated self-responses. While test set performance was high (> 0.9 F1) with and without the 10 kHz tone used, we concluded that the 10kHz tone made ZF-AIM-detector more robust to the differences between its training (which consisted of bird-bird interactions), and the ZF-AIM-bird interactions it was used for.

#### 1.4.3 ZF-AIM-encoder and ZF-AIM-decoder

The call encoder (ZF-AIM-encoder) is used to convert a call waveform to a call token (i.e. an integer), which is the format expected by the event-based interaction model ZF-AIM-interact. The call decoder (ZF-AIM-decoder) is used to convert a call token back to a call waveform.

ZF-AIM-encoder consists of two components: a call acoustic encoder followed by a call token encoder. The call acoustic encoder converts the variable-length call waveform into a variable-length sequence of acoustic feature vectors. The call token encoder converts this variable-length sequence into a single integer. We adopted this two-stage encoding approach because it allowed us to take advantage of recent advances in neural audio codecs (which were used for the call acoustic encoder) before applying further compression (through the call token encoder).

Similarly, ZF-AIM-decoder consists of two components: a call token decoder followed by a call acoustic decoder. Given a call token, the call token decoder autoregressively predicts a sequence of multi-layer acoustic tokens. Then, this sequence is converted to a call waveform by the call acoustic decoder.

##### Call acoustic encoder and decoder

The call acoustic encoder and decoder were the encoder and decoder of an Encodec model. This Encodec was trained with recordings from Pairs 1-34. For the model hyperparameters, we followed closely the setting file, *encodec_base_causal, yaml*, provided by the Audiocraft repository^7^. The only changes are that we used 4 layers of Residual Vector Quantization (RVQ) instead of 32 layers. We observed that 4 layers was enough to yield good reconstruction audio quality for the zebra finch sounds which have relatively low variety (See Table 2 for quantitative results). The model was trained with cropped call audio, including both denoised and non-denoised versions of each call (denoising described above). The model was trained for 50 epochs, using the default training settings from Audiocraft^8^. We observed that for this task, training Encodec with the zebra finch data led to better call reconstruction than using the xeno-canto dataset (which we used for pre-training the call detection model); this is shown in Table 3.

**Table 2:** Reconstruction audio quality (FAD: lower better) and codebook usage for different codebook sizes of the call token encoder-decoder. Measured on Pairs 35-40.

| Codebook size | 32 | 64 | 128 | 256 | 512 |
| --- | --- | --- | --- | --- | --- |
| FAD | 0.486 | 0.506 | 0.399 | 0.444 | 0.431 |
| Codebook usage | 32 | 64 | 128 | 187 | 188 |

**Table 3:**
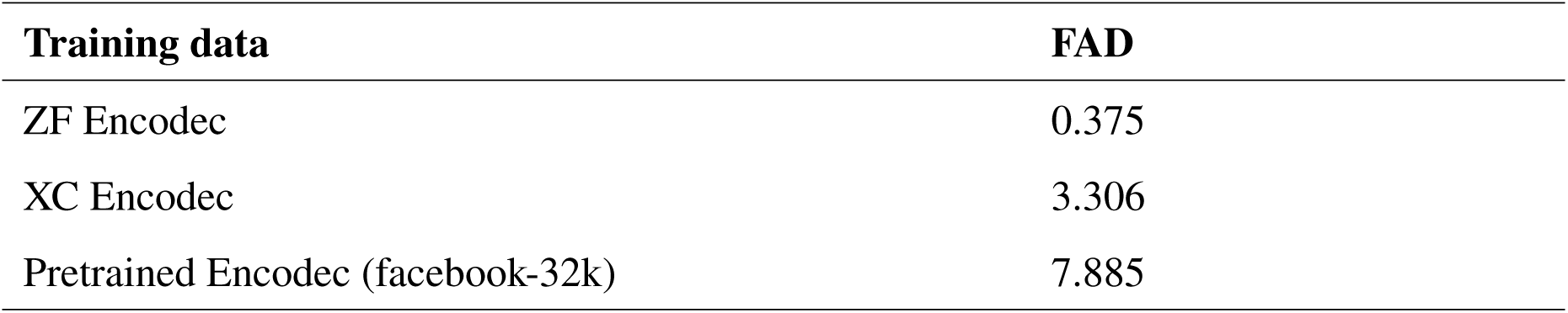
Encodec Reconstruction Audio Quality (FAD: lower better). Pairs 35-40.

The output of the Encodec encoder is a sequence of 128-dim feature vectors at 50 Hz frame rate. Each feature vector is converted into 4 integers (i.e. *multi-layer acoustic tokens*) through the 4 layers of RVQ. Therefore, the bottleneck of the Encodec encoder-decoder has two representations: 1) a sequence of the acoustic feature vectors, and 2) a sequence of the multi-layer acoustic tokens from the RVQ module.

##### Call token encoder and decoder

Given a (non-denoised) call, the input to the call token encoder is the sequence of acoustic feature vectors of that call produced by the call acoustic encoder. The call token encoder is a transformer with recurrent memory followed by a vector quantization layer with a learned codebook. This converts the variable-length sequence of acoustic feature vectors to a single integer, the call token. The call token decoder is a transformer with recurrent memory.

The call token encoder and the call token decoder were trained jointly, as an autoencoder. Training optimized the reconstruction quality of the call token decoder. Given a call, the training target for the output of the call token decoder was the sequence of the multi-layer acoustic tokens produced from the *denoised* version of the same call. These targets were computed during training using the call acoustic encoder, and the weights of the call acoustic encoder were frozen during training. To handle ordering the sequences of the multi-layer acoustic tokens of the denoised call signal in the output, the Delay Pattern from (*62*) was used.

The call token encoder and decoder were trained for 250,000 steps with a batch size of five calls, Adam optimizer, and a learning rate of 1e-4. These hyperparameters were fixed based on results from initial experiments. Training minimized cross-entropy loss, applied to the multi-layer acoustic tokens produced by the call token decoder. The codebook size was tuned as a hyperparameter (described below), with final value of 128. The call token decoder was a transformer with recurrent memory.

We used the sequence of the multi-layer acoustic tokens as the prediction target because it led to a simple classification-based task formulation. In the input, instead of the multi-layer acoustic tokens, we use the acoustic feature vectors to save an extra layer of processing for mapping the multi-layer acoustic tokens to vectors.

Note that, as described above, given a call, the input uses the representation from the un-denoised call signal, while the output predicts the representation from the denoised call signal. This choice, which follows (*63*), was made for the following reasons. On the one hand, the denoising process is computationally expensive, so we want to avoid it during the live interaction. On the other hand, we want the generated call audio to be as clean as possible. Therefore, the model uses the un-denoised signal as the input but predicts the denoised acoustic tokens.

Individual animals often have unique vocal repertoires or signatures (*1*). Therefore, to increase the consistency of the call audio generated by the call decoder across an interaction session, we conditioned the call token decoder on the bird ID in addition to the call token. To accommodate both inputs, the call token and bird ID were each converted to equal-length vectors using a learned embedding layer. These two vectors were summed together and then fed to the subsequent layers in the network. Below, we describe the impact of this individual-specific conditioning on audio quality.

##### Evaluation

ZF-AIM-encoder and ZF-AIM-decoder were evaluated through reconstruction of calls, using Frechet audio distance (FAD) (*64*) ^9^. We first evaluated the reconstruction with the call acoustic encoder and the call acoustic decoder (Encodec encoder-decoder). Three settings were tested: 1) the Pretrained Encodec (facebook-32k) provided by the official Audiocraft repository, 2) an Encodec trained with the xeno-canto dataset, and 3) an Encodec trained with the zebra finch calls in Pairs 1-34. The result is shown in Table 3. The model trained with the zebra finch data achieved the best FAD. This model was used as the call acoustic encoder and the call acoustic decoder in the rest of the paper.

We also evaluated how the vector-quantization codebook sizes of the call token encoder and the call token decoder affect the reconstruction. The one with codebook size 128 achieved the best FAD (0.399) as shown in Table 2 so it was used in all following steps. We also checked the codebook usage of each model. On the validation set, the codebook was fully utilized in the models with size 32, 64, and 128, but not for larger codebooks. Furthermore, audio quality did not always improve as codebook size increased. Finally, audio reconstruction quality when using the entirety of ZF-AIM-encoder plus ZF-AIM-decoder was similar to the reconstruction quality when using the call acoustic encoder-decoder alone (FAD of 0.399 versus 0.375, respectively). This justifies the use of the call token encoder/decoder, as the compression it introduces does not substantially harm performance.

Finally, we evaluated how conditioning the call token decoder on bird ID affected the call reconstruction. We asked, first, whether including bird ID improves audio quality, and second, whether including bird ID makes the reconstructed calls better match the overall qualities of the bird that made the call.

To answer the first question, we compared FAD for call token decoders with and without conditioning on bird ID. Reconstruction quality was similar when conditioning on bird ID (FAD = 0.399) as when not conditioning on bird ID (FAD = 0.355).

To answer the second question, we trained a classifier to predict the bird ID from a call. The architecture and the input of the classifier are exactly the same as the call token encoder. It is a transformer with recurrent memory, and the input is a sequence of acoustic feature vectors produced by the call acoustic encoder. The output is an integer representing a specific bird. The classifier was trained with the calls from Pairs 1-34, and tested with the calls from Pairs 35-40 (the birds in these pairs appeared in the previous 34 pairs). It was trained for 250,000 steps with a mini-batch size of five calls, Adam optimizer, and a learning rate 1e-4. On the test set, the classifier obtained F1 score of 0.419.

We then passed each call in the test set through ZF-AIM-encoder and ZF-AIM-decoder, and evaluated bird ID classification performance on the reconstructed calls. When ZF-AIM-decoder was conditioned on bird ID, the reconstructed call was easier to classify to individual (F1=0.772) than when ZF-AIM-decoder was not conditioned on bird ID (F1=0.081). This confirms that conditioning ZF-AIM-decoder on a bird ID indeed adds individual acoustic signature in the reconstructed audio.

#### 1.4.4 ZF-AIM-interact

Female zebra finches produce short calls that often occur in bursts of activity followed by minutes of silence. To model the interaction between two birds, we treat these calls as discrete, instantaneous events (*CALL* events) separated by periods of silence (*WAIT* events). Following prior work in midi audio generation (*55–57*), we employ a sequence-to-sequence event-based interaction model (ZF-AIM-interact) which predicts the next event based on the sequence of past events.

##### Data Format

For ZF-AIM-interact, inputs and outputs are organized as a sequence of events (Table 4), beginning with a *START* event and then alternating between two types of events, *WAIT* events and *CALL* events.

**Table 4:** Model input and output events.

| Name | Number of events |
| --- | --- |
| *Input events* |  |
| START | 1 |
| WAIT | 24 |
| SELF CALL | 128 |
| PARTNER CALL | 128 |
| NULL CALL | 1 |
| *Output events* |  |
| WAIT-S2S | 24 |
| WAIT-P2S | 24 |
| WAIT-S2P | 24 |
| WAIT-P2P | 24 |
| SELF CALL | 128 |
| PARTNER CALL | 128 |

*WAIT* events represent the duration *dt* in between vocalizations. The duration *dt* is assigned to the closest time in one of 24 discrete categories, referred to as the *dt* categories: 0.01 = 0.01 × 2^0^, 0.02 = 0.01 × 2^1^, 0.04 = 0.01 × 2^2^, 0.01 × 2^23^ (in seconds).

*CALL* events represent the type (self or partner) of the individual who made the vocalization, in addition to the call token of the vocalization (produced by ZF-AIM-encoder). There are therefore 256 *CALL* events, 128 for each of self-calls and partner-calls. There is one additional *CALL* event, a NULL *CALL*, whose use is explained below.

Therefore, input sequences are of the following form:

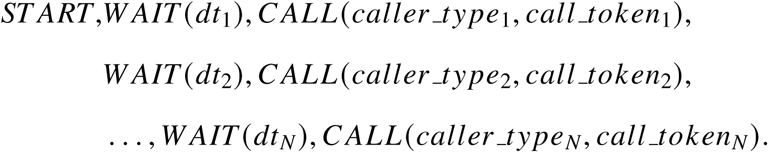

We observed that the duration between vocalizations depends on whether a bird was responding to herself or her partner (Figure 1). Therefore, we designed ZF-AIM-interact to predict the wait time and identity of the responder at the same time. So, for ZF-AIM-interact outputs, *WAIT* events also included the identity (self or partner) of two neighboring *CALLS* events. With this modification, output sequences are of the following form:

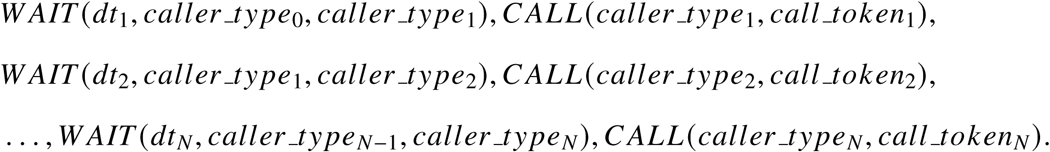

With this addition, there are 96 = 4 × 24 categories of output *WAIT* tokens, one for each of *s2s* (self-to-self), *s2p* (self-to-partner), *p*2*s* (partner-to-self), and *p*2*p* (partner-to-partner).

##### Modeling Task

Given an input sequence of *START*, *WAIT*, and *CALL* events, the task performed by ZF-AIM-interact is to predict the next event in the sequence. Predictions are constrained, so that each *WAIT* event must be followed by a *CALL* event and vice-versa. *WAIT* events following self calls must to be *s2s* or *s2p* and similarly for partner calls. *CALL* events following *s2s* and *ps WAIT* events must be self calls, and similarly for partner calls.

##### Training Data

To construct one training sequence, a start call was sampled randomly from the recorded bird pair interactions; all the information except for the timestamp was removed from this call. This became the *START* event, and its timestamp was used to derive the first *WAIT* event. One of the two bird IDs involved in the interaction was randomly assigned to be self; the other to be the partner. One training sequence consisted of *WAIT* and *CALL* events for the 100 consecutive calls following the *START* event.

We observed that different amount of silence before and after a call waveform could affect the call token derived from the Call Encoder. We adopted the following augmentation process to make ZF-AIM-interact more robust to the amount of silence due to the detection. For each call waveform, we sampled a left silence duration and a right silence duration from [0, 0.2] (seconds), and the call waveform was left-padded and right-padded by zeros with those durations, respectively. This process was done 19 times to derive 20 variants of the call waveform, including the original one that was not padded. Therefore, each call had a set of 20 variant tokens, with potential repetitions. During the training of ZF-AIM-interact, for each call that occurred, one token was sampled uniformly at random from these 20 tokens as a representative.

During training, a partner *CALL* event had a 20% chance to be replaced with a NULL *CALL* event before it was input into ZF-AIM-interact. A NULL *CALL* was used to represent the situation in which ZF-AIM-interact predicted the partner would make the next call, but the partner did not make any call. NULL *CALL* events were not included in the target sequence. Details are given below.

##### Architecture

ZF-AIM-interact is a transformer with recurrent memory. Inputs are *START*, *CALL*, and *WAIT* events, as described above, in addition to metadata: the ID for each bird, the day number of the trial, and the time since light on in the recording chamber. Each of the events and each of these metadata were converted to a vector with a learned embedding layer; these metadata vectors were concatenated with each event vector before being passed into ZF-AIM-interact.

##### Training

ZF-AIM-interact was trained to predict the next event in the sequence, using categorical cross-entropy loss *Loss*_*CE*_ across the possible categories of the next event. As described above, there were 96 possible categories of output *WAIT* events and 256 possible categories of *CALL* events, although at any given time some of these categories were unavailable due to the constraints imposed.

In the preliminary experiments, we observed that ZF-AIM-interact produced much shorter waiting times compared to the distribution from the real bird interactions. To remedy this, we added a KL divergence loss term *Loss*_*KL*_, which is defined as the sum of four terms: 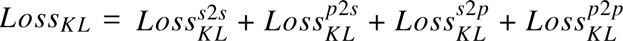, where *p*2*s* corresponds to intervals between a partner call and a self call, and similarly for *p*2*p*, *s2p*, and *s2s*. For one mini-batch seen during training, the term 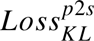 is the negative of the KL divergence between the distribution of wait times for the partner-to-self *WAIT* tokens that are predicted by ZF-AIM-interact for that batch, and the actual distribution of partner-to-self wait times based on the entire dataset. The other terms 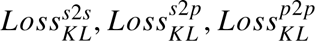 are computed similarly.

ZF-AIM-interact was trained to minimize *Loss*_*CE*_ + *Loss*_*KL*_, the sum of the cross-entropy and KL divergence loss terms.

##### Implementation details

ZF-AIM-interact was trained for 100,000 steps with a mini-batch size of five, Adam optimizer, and a learning rate of 1e-4. It was trained with teacher-forcing (*55, 65*) on entire sequences. Hyperparameters were selected based on results from initial experiments. The batch size of five was the maximum possible due to GPU memory limits.

##### Evaluation

We evaluated the interaction model on the validation set (Pairs 35-40), on three prediction tasks: next caller identity (i.e. partner or self), next call token (i.e. the integer output of the the call encoder), and the wait time until the next call (i.e. the *dt* category). We measured classification accuracy for each of these. For wait time prediction, we additionally measured root mean-squared error (in seconds). As a baseline, we compared to the outputs of a model which made categorical predictions randomly, with probability mass equal to the relative frequencies observed in Pairs 1-34. Results are presented in Table 5. In all metrics, for all predictions, the interaction model performed better than the baseline.

**Table 5:** Prediction metrics for interaction model, compared to random baseline. For accuracy (Acc.), higher scores are better (1 optimal). For root mean-squared error (RMSE), lower scores are better (0 optimal).

| Metric | Model | Baseline |
| --- | --- | --- |
| Caller ID (Acc.) | 0.632 | 0.496 |
| Call Token (Acc.) | 0.056 | 0.010 |
| Wait Time (Acc.) | 0.188 | 0.159 |
| Wait Time (RMSE) | 976.0 | 1126.4 |

As described above, a KL divergence loss (*Loss*_*KL*_) was used in addition to the main cross-entropy loss (*Loss*_*CE*_). The *Loss*_*CE*_-only model achieves *Loss*_*CE*_ = 3.329 and *Loss*_*KL*_ = 0.333 on the validation set, while the model with both loss terms achieves a *Loss*_*CE*_ = 3.351 and a *Loss*_*KL*_ = 0.0624. Therefore, *Loss*_*KL*_ is improved greatly on the validation set when the *Loss*_*KL*_ is optimized during training, with only a minor change in *Loss*_*CE*_..

In Fig. 6, we compare the wait time distribution from the two-bird interactions, with the wait time distribution from the *in silico* interactions between two copies of ZF-AIM. The inclusion of the KL divergence loss makes the prediction and the real probabilities closer, especially for the overly-predicted categories such as 0.16-sec and 0.32-sec in partner-to-self *WAIT* events.

**Figure 6:**
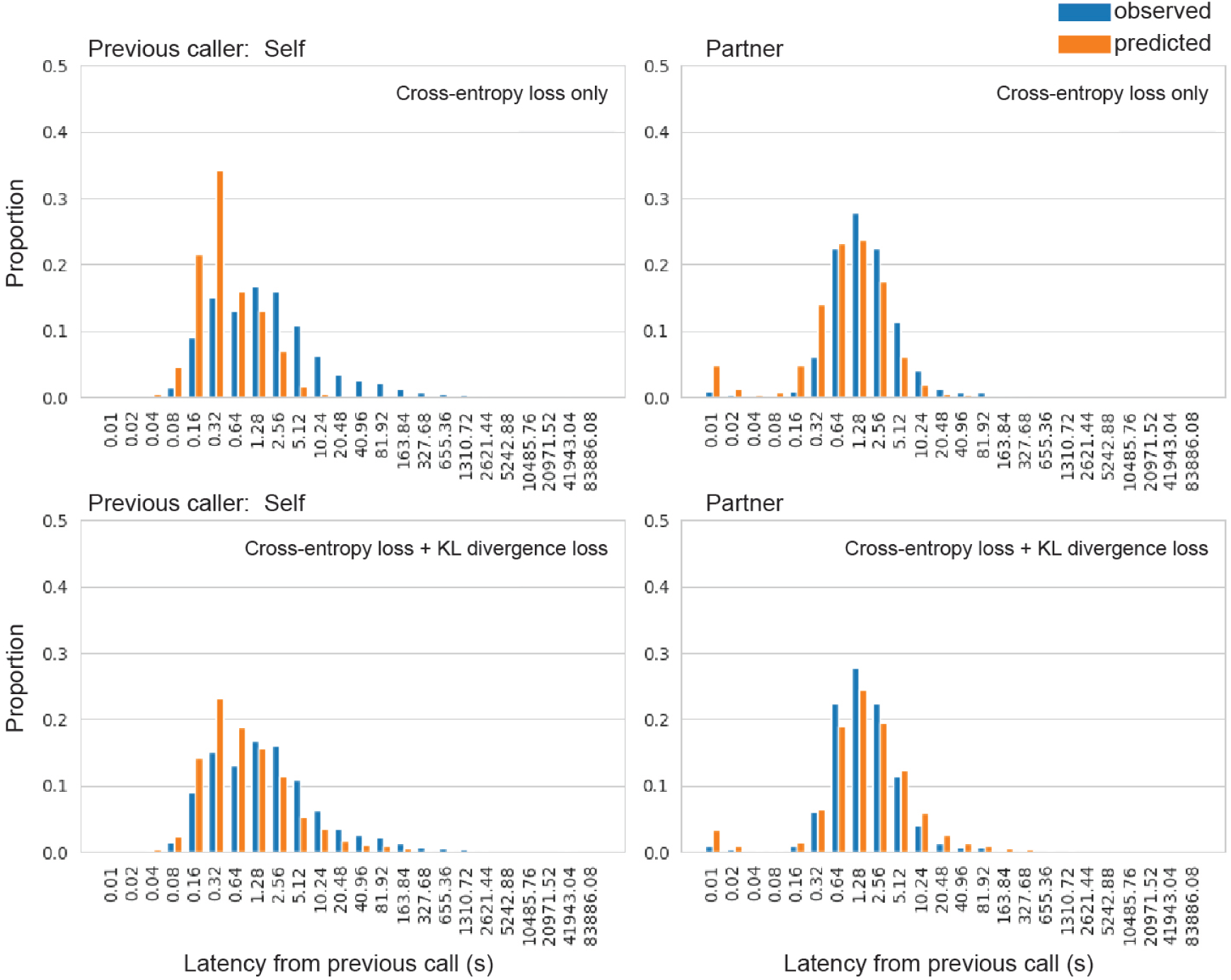
The effect of the KL divergence loss on the wait time prediction. Panels: (top) cross-entropy loss only, (bottom) cross-entropy loss + KL divergence loss.

### 1.5 Interaction

We use ZF-AIM to interact with a partner, which can be either a zebra finch or another copy of ZF-AIM. To do so, a ‘self’ bird id and a ‘partner’ bird id are chosen and provided to ZF-AIM during the interaction. During an interaction session, as calls occurred, ZF-AIM predicts the identity of the next caller, the call token of the next call, and the wait time until the next call.

Before beginning the interaction session, an empty event history memory buffer is initialized. In what follows, when an event is referred to as being *processed*, it means that the event is input into to ZF-AIM-interact and the event is added to the event history memory buffer.

At the outset of an interaction session, ZF-AIM-interact is provided with a *START* event with the start timestamp *ts*_0_. This event is processed and ZF-AIM-interact predicts a *WAIT* event (i.e. it predicts the caller ID of the next call and wait time until the next call). Then, a *checkpoint* is created at time *ts*_1_ = *ts*_0_ + *dt*. If this checkpoint is reached and no more events have occurred, there are two possibilities: 1) the predicted *caller_type* was self (ZF-AIM), or 2) the predicted *caller_type* was the partner. Alternatively, an event might occur before the checkpoint was reached. We now describe the behavior in each of these scenarios.

Assume the checkpoint is reached and the predicted *caller_type* was self. Then, the predicted *WAIT* event is processed, ZF-AIM-interact predicts a *CALL* event, ZF-AIM-decoder generates call audio, and the *CALL* event is processed. Finally, a new *WAIT* event (i.e. a new wait time *dt* and a new *caller_type*) is predicted by ZF-AIM-interact. A new checkpoint, at time *ts_2_ = ts_1_ + dt*, is produced based on *dt* and the process goes on.

Assume the checkpoint is reached and the predicted *caller_type* was partner. Then, the *WAIT* event is processed, and a NULL *CALL* event is processed. Finally, a new *WAIT* event (i.e. a new wait time *dt* and a new *caller_type*) is predicted by ZF-AIM-interact. A new checkpoint, at time *ts*_2_ = *ts*_1_ + *dt*, is produced based on *dt* and the process goes on. In other words, if ZF-AIM reaches a checkpoint with the predicted next caller being the partner, it means that the partner hasn’t actually called (otherwise, this checkpoint would have been interrupted, which is explained in the next paragraph). Therefore, we used a NULL *CALL* event to represent this situation.

The process described so far assumes that there were no partner *CALL* events before the checkpoint was reached. What if there was a new partner call at **ts*′_1_* before the checkpoint at *ts*_1_ was reached? Then, the predicted *WAIT* event is *not* processed. Instead, a *WAIT* event is created based on the timestamp of the partner call and the last event in the event history, *WAIT* (*ts*′_1_ − *ts*_0_), and this new *WAIT* event is processed. Then, the partner *CALL* event would be processed, and a new *WAIT* event (i.e. a new wait time *dt* and a new *caller_type*) is predicted by ZF-AIM-interact. A new checkpoint, at time 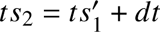, is produced based on *dt* and the process goes on.

In an interaction, to avoid a situation where the wait times between calls played by ZF-AIM were a limited set of discrete values, wait times between events are randomly adjusted. After ZF-AIM-interact predicts a *dt* category *C* for a *WAIT* event, if the *caller_type* of the wait event was self, then the next self call would be played at a time randomly drawn from the interval [(*C* + *L*)/2, (*C* + *R*)/2] where *L* was the next-smallest *dt* category to *C* and *R* was the next-largest *dt* category to *C*. For example, if the *dt* category *C* = 0.01-sec is predicted, then a wait time was sampled from [(0 + 0.01)/2, (0.01 + 0.02)/2] = [0.005, 0.015]. If the *dt* category *C* = 0.02-sec is predicted, then a wait time was sampled from [(0.01 + 0.02)/2, (0.02 + 0.04)/2] = [0.015, 0.03].

### 1.6 Settings for live interactive playbacks

We used sounddevice ^10^ to handle streaming input and output. Audio is streamed using sounddevice in chunks with duration 0.02 seconds at a sampling rate of 32 kHz, and then collected into 0.1-second chunks before being passed to ZF-AIM-detector. Sounds produced by ZF-AIM were played through a speaker.

To save processing time during the live interaction for converting a call token to a call waveform, we pre-generated 1000 waveforms by using different random seeds for each call token, using ZF-AIM-decoder. There were 128 unique call tokens, so this amounts to 128 × 1000 = 128, 000 pre-generated waveforms for each bird. During the live interaction, when ZF-AIM-interact predicted a call token, one call waveform was sampled at random from the 1000 call waveforms produced with that token.

### 1.7 Analysis

#### 1.7.1 Data selection

We divided the data into four 3.5 hour time bins per day, starting from lights on. Because birds were paired on the afternoon of day 1 and recorded for 48 hours, we included the final time bin of day 1, all four time bins from day 2, and the first two time bins of day 3 for a total of 7 analysis bins (Fig. 1B). Our final dataset for analysis consisted of 834,309 calls produced by 21 birds across 40 pairwise live interactions. From passive playbacks, we recorded 222,933 calls produced by 14 birds across 30 playback sessions (most of the birds were recorded with both fixed-interval passive playback and random-interval passive playback; moreover, 4 birds were tested on 2 fixed-interval passive playback sessions and 5 birds were tested on 2 random-interval passive playback sessions). From interactive playbacks, we recorded 144,349 calls produced by 11 birds across 22 playback sessions (all birds were recorded with both ZF-AIM and ZF-AIM-ablated), as well as 111,456 calls produced by the playback models during those sessions.

To analyze how birds modulated their calling behavior during rapid interactions, we categorized calls as “direct responses” if the call started within 1 second from the end of the partner call (whether the partner was a bird or a playback). Most other calls produced by the bird were considered “other calls”. However, if a bird produced a string of calls with less than 1 second of silence between the calls (i.e., direct responses to self), only the first was considered in these analyses as either a “direct response” or “other call” depending on what preceded the first call. We similarly categorized all calls produced by a bird’s partner (whether a live bird or a playback) as “directly responded to” or “other” using similar logic (Fig. 7 for a schematic of these definitions). All of the selection criteria described here were preregistered^11^ prior to analyzing any of the interactive playback data included in the study.

**Figure 7:**
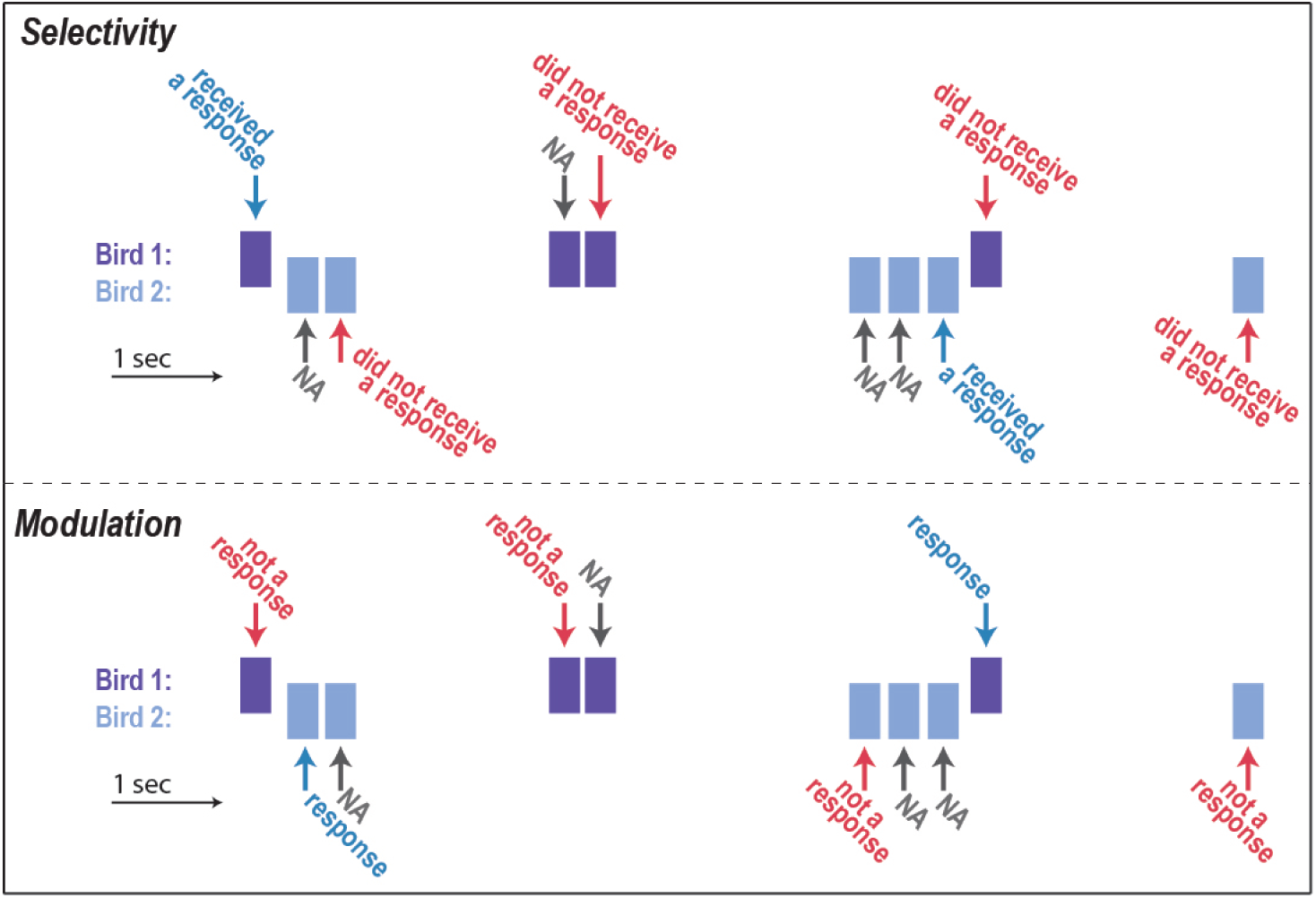
Diagram outlining logic for defining calls that receive a response and calls that are responses. Each colored rectangle indicates a simulated call from bird 1 (purple) or bird 2 (blue) in a theoretical pair, with time on the x-axis. Top diagram indicates the logic for determining which calls receive a response. Calls followed by a partner call at low latency are considered as having received a response, whereas calls with no immediate call following are not (did not receive a response). However, calls that are followed by a rapid response from the same bird are not considered in the analysis (NA). Similarly, the bottom diagram indicates the logic for determining which calls are responses to the partner. Specifically, a call that occurs rapidly after the partner is considered a response, whereas a call that is not produced rapidly after another call is considered not a response. However, if a bird produces a rapid call after itself, the call is not considered in the analysis (NA). We note that this logic was preregistered prior to analyzing bird interactions with ZF-AIM, and that the excluded calls do not meaningfully change any analysis (rapid self responses are rare; see Fig. 1).

#### 1.7.2 Acoustic features

To assess vocal flexibility, we measured acoustic features of calls (see above). We then performed a principal components analysis (PCA; all features centered and *z*-transformed prior). We focused our analyses on PC1 and PC2, while also including supplementary analyses on three features selected^12^*a priori*: duration, entropy, dominant frequency (DF). The loadings of each acoustic feature on each PC are in Table 6.

**Table 6:**
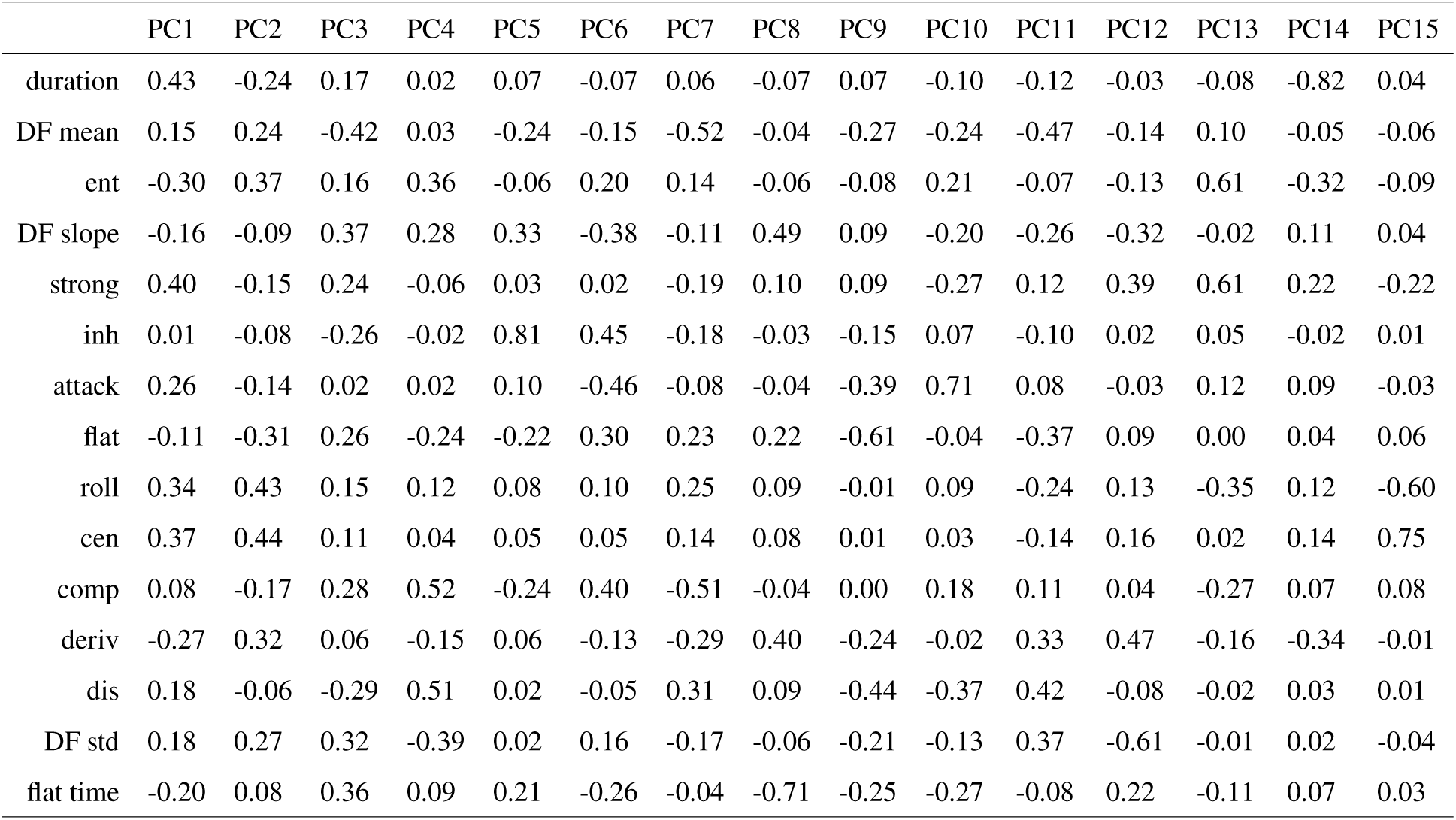
Loadings for each acoustic feature for all principal components.

|  | PC1 | PC2 | PC3 | PC4 | PC5 | PC6 | PC7 | PC8 | PC9 | PC10 | PC11 | PC12 | PC13 | PC14 | PC15 |
| --- | --- | --- | --- | --- | --- | --- | --- | --- | --- | --- | --- | --- | --- | --- | --- |
| duration | 0.43 | -0.24 | 0.17 | 0.02 | 0.07 | -0.07 | 0.06 | -0.07 | 0.07 | -0.10 | -0.12 | -0.03 | -0.08 | -0.82 | 0.04 |
| DF mean | 0.15 | 0.24 | -0.42 | 0.03 | -0.24 | -0.15 | -0.52 | -0.04 | -0.27 | -0.24 | -0.47 | -0.14 | 0.10 | -0.05 | -0.06 |
| ent | -0.30 | 0.37 | 0.16 | 0.36 | -0.06 | 0.20 | 0.14 | -0.06 | -0.08 | 0.21 | -0.07 | -0.13 | 0.61 | -0.32 | -0.09 |
| DF slope | -0.16 | -0.09 | 0.37 | 0.28 | 0.33 | -0.38 | -0.11 | 0.49 | 0.09 | -0.20 | -0.26 | -0.32 | -0.02 | 0.11 | 0.04 |
| strong | 0.40 | -0.15 | 0.24 | -0.06 | 0.03 | 0.02 | -0.19 | 0.10 | 0.09 | -0.27 | 0.12 | 0.39 | 0.61 | 0.22 | -0.22 |
| inh | 0.01 | -0.08 | -0.26 | -0.02 | 0.81 | 0.45 | -0.18 | -0.03 | -0.15 | 0.07 | -0.10 | 0.02 | 0.05 | -0.02 | 0.01 |
| attack | 0.26 | -0.14 | 0.02 | 0.02 | 0.10 | -0.46 | -0.08 | -0.04 | -0.39 | 0.71 | 0.08 | -0.03 | 0.12 | 0.09 | -0.03 |
| flat | -0.11 | -0.31 | 0.26 | -0.24 | -0.22 | 0.30 | 0.23 | 0.22 | -0.61 | -0.04 | -0.37 | 0.09 | 0.00 | 0.04 | 0.06 |
| roll | 0.34 | 0.43 | 0.15 | 0.12 | 0.08 | 0.10 | 0.25 | 0.09 | -0.01 | 0.09 | -0.24 | 0.13 | -0.35 | 0.12 | -0.60 |
| cen | 0.37 | 0.44 | 0.11 | 0.04 | 0.05 | 0.05 | 0.14 | 0.08 | 0.01 | 0.03 | -0.14 | 0.16 | 0.02 | 0.14 | 0.75 |
| comp | 0.08 | -0.17 | 0.28 | 0.52 | -0.24 | 0.40 | -0.51 | -0.04 | 0.00 | 0.18 | 0.11 | 0.04 | -0.27 | 0.07 | 0.08 |
| deriv | -0.27 | 0.32 | 0.06 | -0.15 | 0.06 | -0.13 | -0.29 | 0.40 | -0.24 | -0.02 | 0.33 | 0.47 | -0.16 | -0.34 | -0.01 |
| dis | 0.18 | -0.06 | -0.29 | 0.51 | 0.02 | -0.05 | 0.31 | 0.09 | -0.44 | -0.37 | 0.42 | -0.08 | -0.02 | 0.03 | 0.01 |
| DF std | 0.18 | 0.27 | 0.32 | -0.39 | 0.02 | 0.16 | -0.17 | -0.06 | -0.21 | -0.13 | 0.37 | -0.61 | -0.01 | 0.02 | -0.04 |
| flat time | -0.20 | 0.08 | 0.36 | 0.09 | 0.21 | -0.26 | -0.04 | -0.71 | -0.25 | -0.27 | -0.08 | 0.22 | -0.11 | 0.07 | 0.03 |

#### 1.7.3 Statistics

To measure correlations in overall calling behavior within a pair, we counted the number of calls produced by each bird and/or playback across 10 min bins. We then conducted linear mixed-effects models with the number of calls from the Right as the dependent variable and the number of calls from the Left as the independent variable. We included 3 random effects: Left Bird ID, Right Bird ID and Pair ID. For measures of overall calling and response rates, we used mixed-effects models with time (numeric variable from 0 (day 1 evening) to 6 (day 3 mid-morning)) as a fixed effect and caller ID as a random effect. To measure these effects across conditions, we conducted full factorial models with condition (interactive partner; e.g., live bird, fixed-interval passive playback, ZF-AIM, etc.) as an additional fixed effect. To measure differences in response latencies across conditions, we calculated the median response latency for all calls produced within at least 4 seconds of the partner and conducted linear mixed-effects models with latency as the dependent variable, condition as the independent variable, and caller ID as a random effect.

For measures of selectivity we calculated mean feature (PC) values for calls that received a response compared to calls that did not receive a response across each entire recording. We then conducted linear mixed-effects models with call response type (responded to vs. not responded to) as the independent variable, an acoustic feature as the dependent variable, and partner ID as a random effect. To compare across pairs of conditions (e.g., between live and *in-silico* interactions), we included condition as a second independent variable in a full factorial model, asking whether the interaction between condition and call response type predicted acoustic features. To directly compare the magnitude of the effects across conditions, we also calculated mean differences in features (e.g., mean feature of all calls that received a response minus the mean feature of all calls that did not receive a response), and conducted linear mixed effects models with the difference as the dependent variable, condition as an independent variable, and partner ID as a random effect. To compare these differences to chance, we conducted intercept-only linear mixed effects models. We conducted similar models for feature modulation, except with the call response type being divided into calls that were produced in response to the partner vs. calls that were not responses to the partner. Caller ID was used as the random effect in all models.

For feature covariation, we subsetted out all calls produced as a response to the partner for each bird in every recording. We then conducted a linear correlation between the acoustic features of these calls and the calls from the partner that they were responding to, and calculated the correlation coefficient. To test whether feature covariation was different from chance, we conducted intercept-only linear mixed effects models with caller ID as a random effect. To compare across conditions, we used similar linear mixed-effects models as described above, with the correlation coefficient as the dependent variable, condition as the independent variable, and caller ID as a random effect.

Statistical analyses were conducted using R 4.2.1. All mixed-effects models were conducted using the lme4 package (*66*) and Tukey’s post-hoc contrasts were calculated using the emmeans package (*67*) after all significant main effects across more than two conditions. For all comparisons of birds in live interactions to playbacks, we limited our live interaction dataset to those birds that were in the playback conditions. The primary analyses presented and statistical framework were designed and preregistered^13^ prior to any analysis of the interactive playbacks in this study.

## Supplementary text

### Live interactions between two birds: analysis of individual acoustic features

In addition to the analyses of vocal flexibility based on variation in principal component space, we also analyzed variation in three acoustic features that are commonly analyzed for vocal communication: duration, spectral entropy, and dominant frequency (DF; see Methods). Selectivity: Calls that received direct responses were longer and lower in entropy than other calls (duration: *F*_1,89.0_ = 34.8, *p* < 0.001; entropy: *F*_1,94.6_ = 38.0, *p* < 0.001; DF: *F*_1,80.3_ = 1.0, *p* = 0.321; Fig. S1A-C). Modulation: Calls that were produced as direct responses to the partner were longer, lower in entropy, and higher in dominant frequency than other calls (duration: *F*_1,87.7_ = 25.8, *p* < 0.001; entropy: *F*_1,96.5_ = 17.2, *p* < 0.001; DF: *F*_1,98.6_ = 9.2, *p* = 0.003; Fig. S1D-F). Response covariation: Across birds, all three acoustic features demonstrated significantly positive correlation coefficients when comparing responses to the calls they were responding to on a response-by-response basis (duration: *t*_19.1_ = 11.8, *p* < 0.001; entropy: *t*_18.9_ = 6.1, *p* < 0.001; DF: *t*_79.0_ = 7.3, *p* < 0.001; Fig. S1G,H). Correlation coefficients were significantly different across features (*F*_1,158.0_ = 108.6, *p* < 0.001), with post-hoc contrasts indicating significant differences for all three pairwise comparisons (*p* < 0.001 for all).

### Live interactions between two birds: analysis within a single putative call “type”

It is possible that observed acoustic flexibility (selectivity, modulation, and covariation) could be primarily driven by the use of distinct call types. For example, birds may selectively respond to only a specific call type. We found evidence against this possibility here. Specifically, we limited our acoustic analyses to a single putative call “type” and continued to find acoustic flexibility.

We trained a classifier on adult zebra finch calls with call type identity (see Methods; (*24,32,33*)) and applied it to our data. The most commonly classified call type was the “distance call” (DC), which constituted on average ≈ 56% of calls produced (Fig. S2A,B). These calls are typically long harmonic stacks, and visual examination of a sample confirmed this. Birds selectively responded to DCs that were higher on PC1 (*F*_1,87.2_ = 34.7, *p* < 0.001; Fig. S2C), with no selectivity on PC2 (*F*_1,92.1_ = 0.3, *p* = 0.580). Birds also modulated the structure of their DCs, where DCs that were in direct response to the partner had higher values for both PC1 and PC2 than DCs that were not direct responses (PC1: *F*_1,92.9_ = 76.3, *p* < 0.0010; PC2: *F*_1,97.7_ = 5.4, *p* = 0.022; Fig. S2D). Finally, the structure of DCs that were in direct response to the partner exhibited covariation with the preceding call for both PC1 and PC2 (PC1: *t*_19.9_ = 14.5, *p* < 0.001; PC2: *t*_15.4_ = 3.8, *p* = 0.002; Fig. S2E). This documentation of selectivity, modulation, and covariation within a call type suggests that our results are not driven simply by the use of different call “types”.

**Figure S1:**
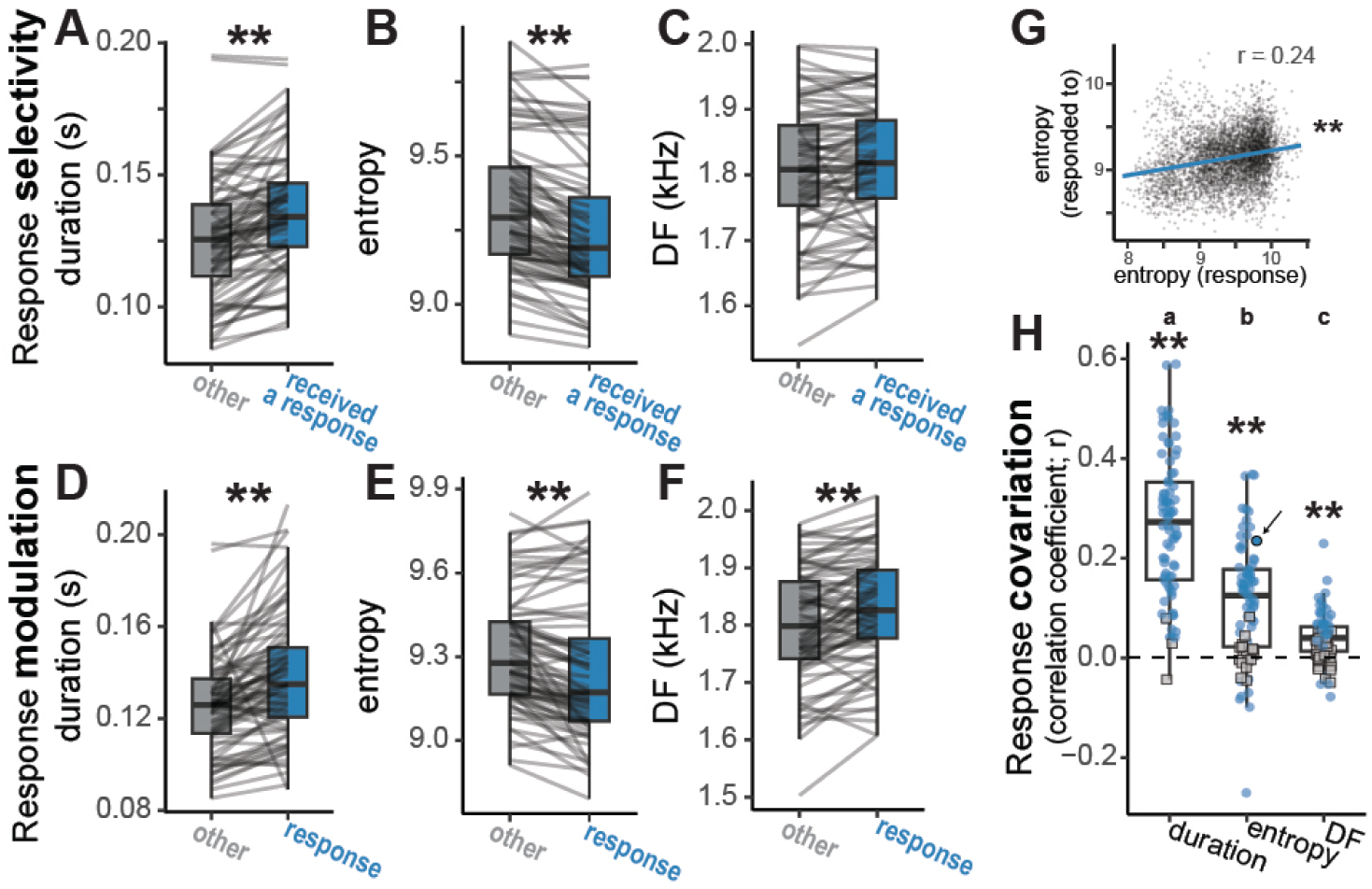
Vocal flexibility based on three acoustic features: duration, entropy and dominant frequency (DF). A-C: Birds selectively respond to calls that are longer (A) and lower in entropy (B). D-F: Birds respond with longer (D), lower entropy (E), and higher DF calls (F). G. An example correlation between the entropy of each response and the entropy of the call being responded to. H. Correlation coefficients between calls and their responses for each acoustic feature. The arrow indicates the correlation depicted in G. Letters above indicate significant differences across features. For all panels, ∗∗ *p* < 0.005.

### Null dataset by swapping pairs

We created a synthetic dataset of 14 pairs in which we “rotated” the left birds from pairs 27-40 forward two positions (e.g., left bird in pair 28 paired with the right bird of pair 30). We calculated the same measures of the interaction on these pairs, as well as compared the magnitude of these effects to the actual data from those pairs (Fig. S3A).

We found a weak but significant correlation in calling across bins (LMM: *F*_1,1421.9_ = 4.2, *p* = 0.040; Fig. S3B), which relates to increases in call rates over time (which is unchanged by our swapping procedure; *F*_1,162.1_ = 11.1, *p* < 0.001; Fig. S3C), and this correlation in call numbers is far weaker than in the natural pairs (*F*_1,2876.0_ = 274.6, *p* < 0.001).

**Figure S2:**
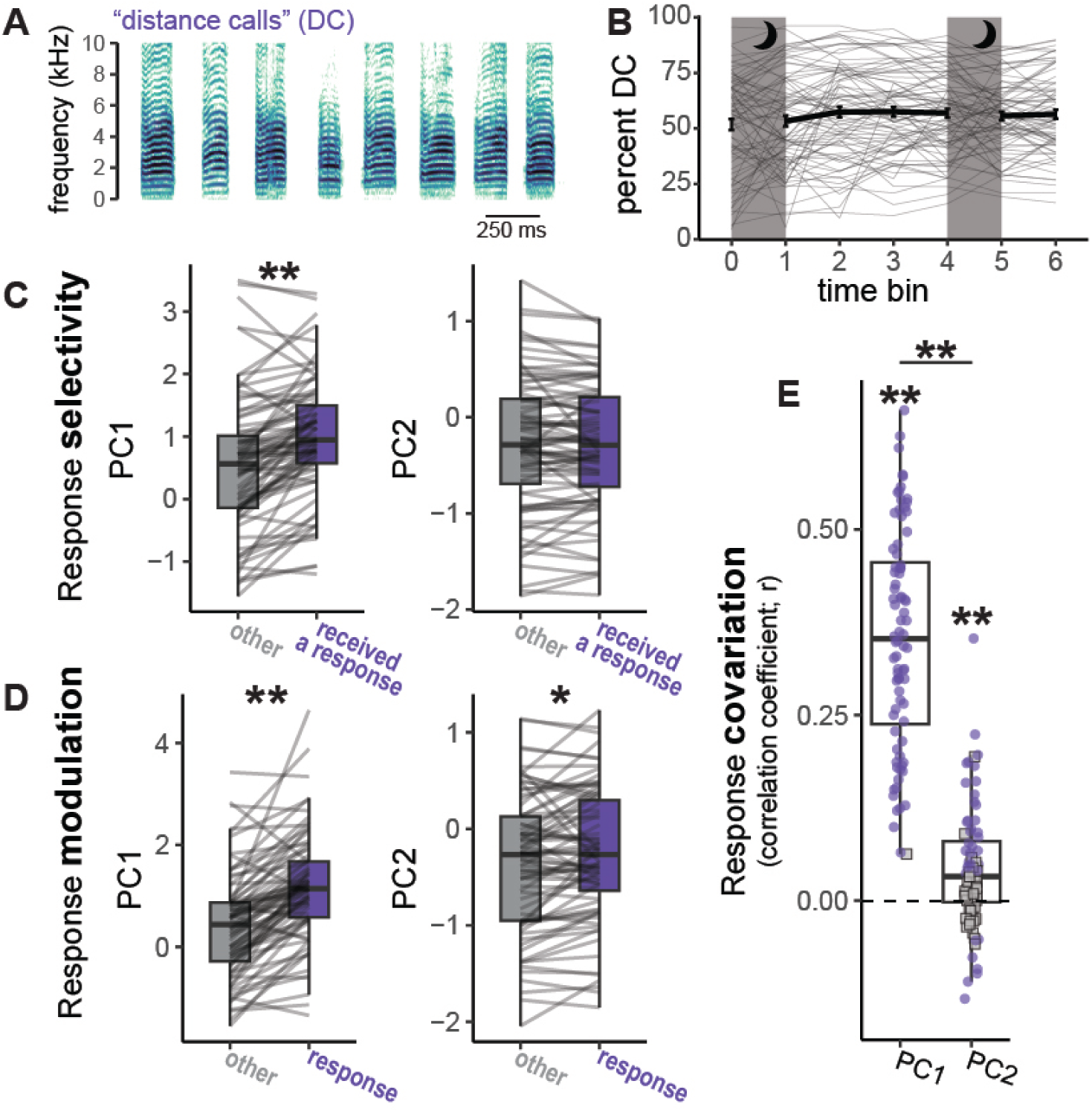
Vocal flexibility analyzed on a single call “type”. A. Examples of distance calls (DCs) (based on (*24*)). B. Percent of calls classified as DCs across time. Each thin line depicts the values from a single bird within a pair, and the thick line depicts the mean ± SE across birds. C. Response selectivity within DCs based on PC1 (left) and PC2 (right). D. Response modulation within DCs based on PC1 (left) and PC2 (right). E. Covariation of DCs used as responses to the call they are in response to for each PC. For C and D, lines connect mean values for a single bird within a pair. Purple dots depict significant correlations for a bird in a pair, while gray squares depict non significant correlations. For all panels, ∗*p* < 0.05, ∗ ∗ *p* < 0.005.

**Figure S3:**
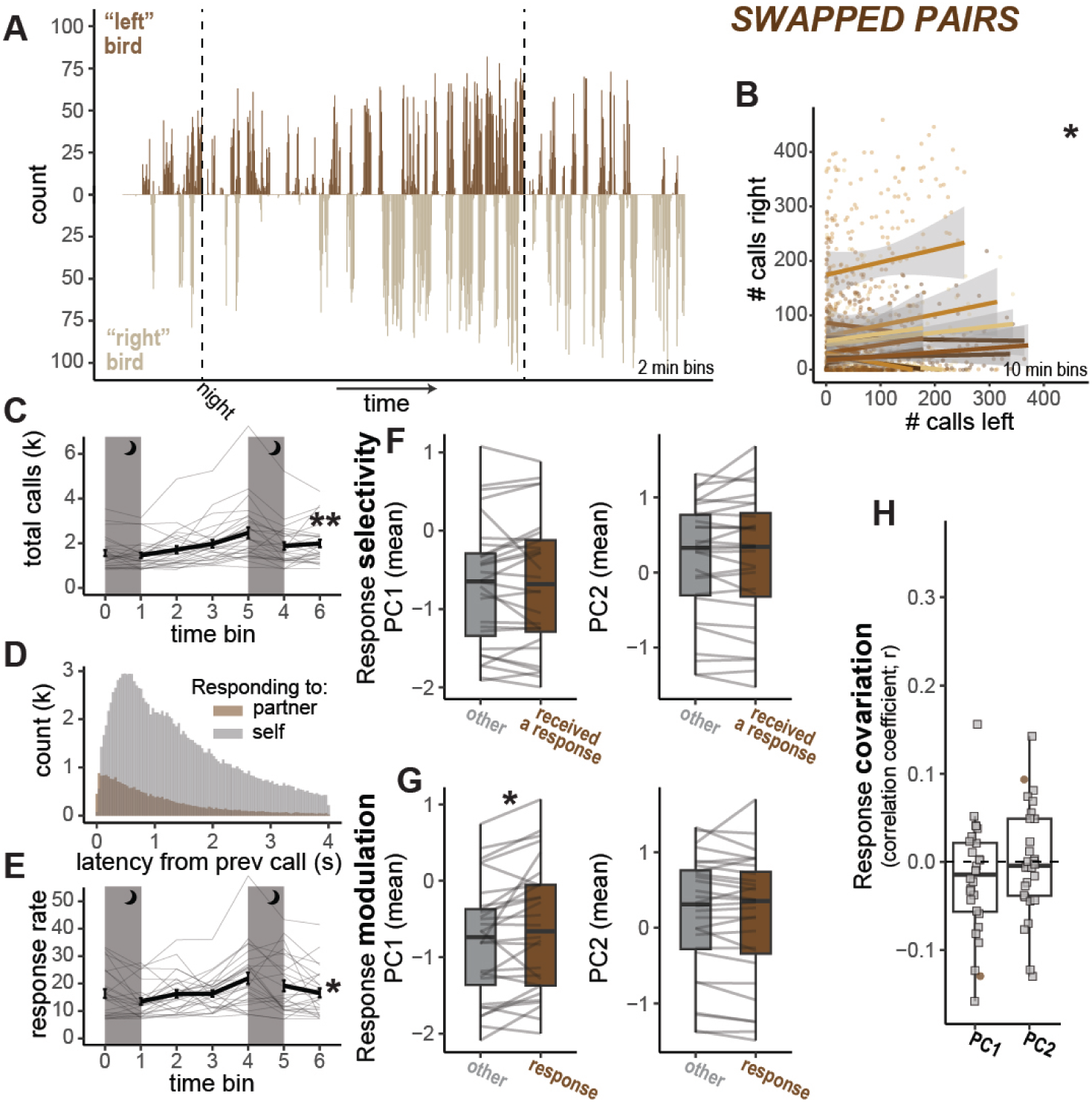
Limited measures of flexibility in channel swapped analysis. A. Example histogram of a channel swapped pair, with the number of calls produced by the left bird (up) and right bird (down) across 2 minute bins. B. Correlations for the number of calls produced in 10 minute bins for each channel swapped pair. Each dot depicts the value for a single bin within a pair. Each line depicts the simple linear correlation for a pair with shading indicating the standard error. C. Total number of calls within each large time bin. D. Response latencies based on whether the previous call was the partner (brown) or the same bird (gray). E. Response rates (percent of calls that receive a response from the partner within 1 second) over time. For C and E, the thin lines depict values from a single bird within a pair, and the thicker line depicts the mean ± across birds. F. Response selectivity based on PC1 (left) and PC2 (right). G. Response modulation based on PC1 (left) and PC2 (right). For F and G, lines depict values for a single bird within a pair. H. Response covariation, with brown circles depicting significant correlations and gray squares depicting non-significant correlations for a given bird in a pair. For all panels, ∗*p* < 0.05, ∗ ∗ *p* < 0.005.

Rapid responses to the partner birds in the swapped pair dataset were rare (≈ 10%; lower than in natural pairs: *F*_1,37.6_ = 102.0, *p* < 0.001; Fig. S3D) with a significant increase over time (*F*_1,166.0_ = 7.3, *p* = 0.008; Fig. S3E). Call properties did not significantly predict whether a call received a response (PC1: *F*_1,27.4_ = 0.0, *p* = 0.992; PC2: *F*_1,27.5_ = 0.2, *p* = 0.630; Fig. S3F), which was significantly different from the live pairs for PC1 (PC1: *F*_1,58.2_ = 23.6, *p* < 0.001; PC2: *F*_1,61.6_ = 1.3, *p* = 0.261).

Interestingly, calls that were classified as direct responses to the partner seemed to be modulated on PC1 (*F*_1,23.6_ = 6.6, *p* = 0.017), but not PC2 (*F*_1,27.4_ = 0.6, *p* = 0.441; S3G). The significant effect for PC1 likely relates to the fact that duration loads heavily on PC1 and the longest calls were most likely to fall within the 1 second response window. However, the magnitude of modulation for both PCs was smaller than in the actual pairs (PC1: *F*_1,63.9_ = 29.4, *p* < 0.001; PC2: *F*_1,64.3_ = 6.4, *p* = 0.014).

Finally, we found no consistent direction of correlation coefficients when analyzing responses and the calls they were in response to for either PC (PC1: *t*_27_ = −1.7, *p* = 0.11; PC2: *t*_13.9_ = 0.0, *p* = 0.963; Fig.S3H). Moreover, the correlations were weaker than in the actual pairs for PC1, and marginally weaker for PC2 when compared to live interactions (PC1: *F*_1,43_ = 279.2, *p* < 0.001; PC2: *F*_1,38.3_ = 3.1, *p* = 0.084).

### ZF-AIM in-silico results

To test whether ZF-AIM could simulate an extended interaction, we reproduced the full dataset of 40 pairs (using the same Bird IDs in each pair). Pairs of *in-silico* models produced correlated numbers of calls (LMM: *F*_1,5415_ = 4322.8, *p* < 0.001; Fig. S4;A,B), with no consistent change in call numbers over time (*F*_1,36.7_ = 1.7, *p* = 0.205; S4C). Across models, ≈ 31% of calls received direct responses, with this rate slightly increasing over time (*F*_1,490.6_ = 6.8, *p* = 0.010; S4D,E). Models were selective in their responses based on PC1 but not PC2 (PC1: *F*_1,95.6_ = 20.4, *p* < 0.001; PC2: *F*_1,96.4_ = 0.1, *p* = 0.738; S4F). Models also modulated their responses based on PC1 but not PC2 (PC1: *F*_1,96.4_ = 34.3, *p* < 0.001; PC2: *F*_1,96.4_ = 0.1, *p* = 0.795; S4G). Finally, models exhibited covariation of their responses for both PCs (PC1: *t*_17.9_ = 20.0, *p* < 0.001; PC2: *t*_16.2_ = 8.9; S4H) with the strength of covariation being different between PCs (*p* < 0.001). While the direction of effects were generally consistent between the *in-silico* agents and between live birds, there were some differences in the magnitude of effects.

**Figure S4:**
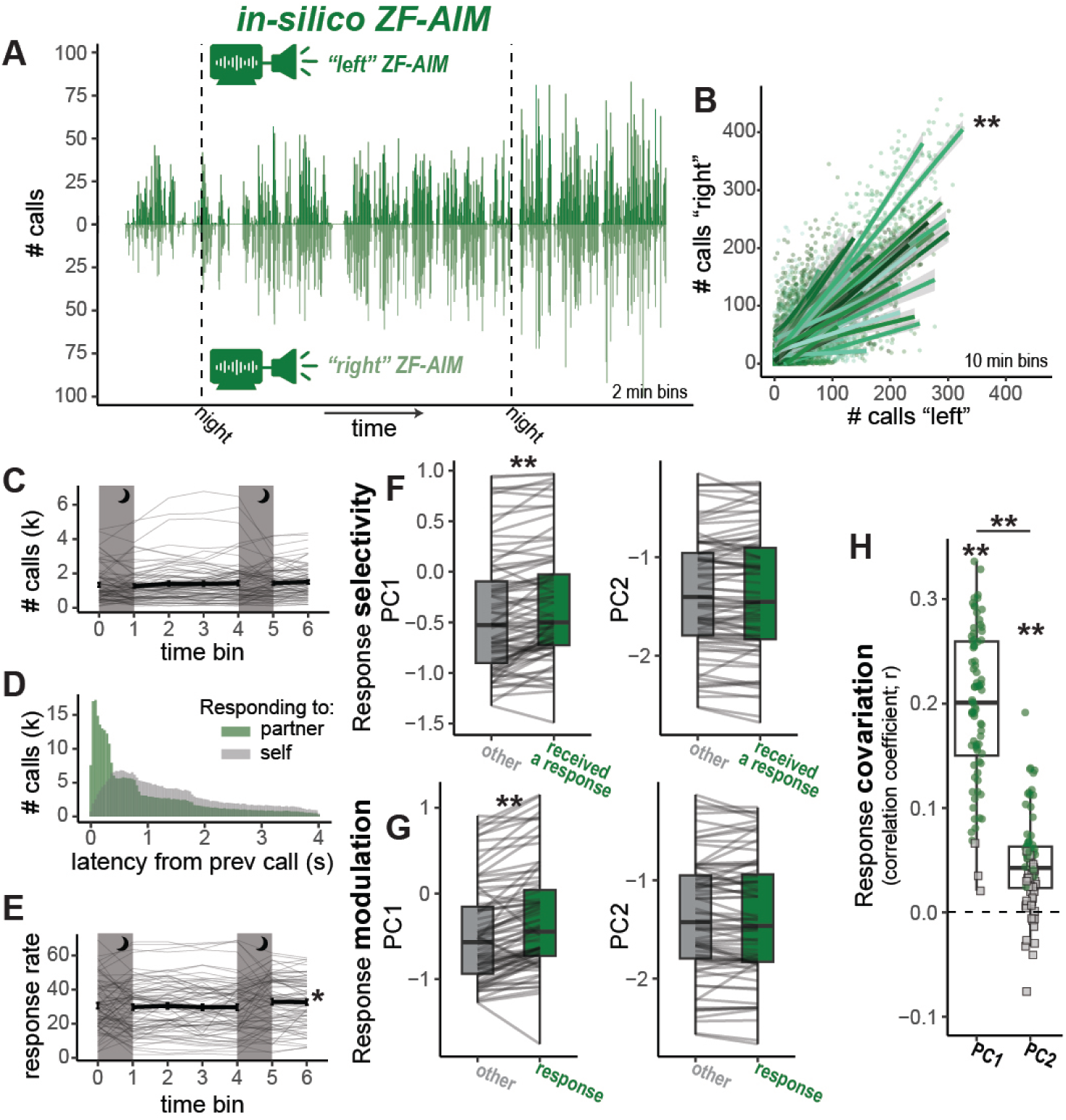
Behavior of ZF-AIM during in-silico interactions with another ZF-AIM. A. Example histogram of calling behavior across 2 min bins. B. Correlations in the number of calls produced across 10 min birds by each agent within a pair. C. Total calls across time. D. Response latencies based on whether the previous call was produced by the partner (teal) or the same agent (gray). E. Response rates over time. For C and E, thin lines depict the behavior of an agent in a pair, and the thick line depicts the mean ± SE across agents. F. Response selectivity based on PC1 (left) and PC2 (right). G. Response modulation based on PC1 (left) and PC2 (right). For F and G, each line depicts the mean value from one agent within a pair. H. Response covariation as measured by correlation coefficients of feature values between calls and their responses. Teal dots depict significant covariation by an agent in a pair, while gray squares depict non-significant covariation. For all panels, ∗*p* < 0.05, ∗ ∗ *p* < 0.005.

The *in-silico* models exhibited tighter correlations in call rates than live birds (*F*_1,9965.8_ = 74.5, *p* < 0.001) but there was no significant difference in the overall change in calling over time between *in-silico* models and live birds (*F*_1,69.3_ = 2.2, *p* = 0.145).

Response rates were generally higher among live birds (*F*_1,85.8_ = 18.6, *p* < 0.001), but there was a similar magnitude of change over time (*F*_1,994.8_ = 0.3, *p* = 0.586). The magnitude of response selectivity *in-silico* was significantly weaker for PC1 (*F*_1,199.2_ = 20.0, *p* < 0.001) and marginally weaker for PC2 (*F*_1,199.2_ = 2.8, *p* = 0.094). Similarly, modulation was weaker *in-silico* relative to live interactions for both PCs: (PC1: *F*_1,218.5_ = 30.6, *p* < 0.001; PC2: *F*_1,214.9_ = 4.4, *p* = 0.036). Covariation *in-silico* was weaker than in live interactions for PC1 (*F*_1,138.2_ = 81.0, *p* < 0.001) but not PC2 (*F*_1,135.2_ = 0.2, *p* = 0.664).

### Ablated model

We analyzed the behavior of ZF-AIM-ablated during live interactions with a bird (Fig. S5A) to confirm our intended effects. The model was designed to eliminate all aspects of acoustic flexibility (selectivity, modulation, and covariation) by removing call acoustic features from the detected calls of the partner bird and by selecting a call for the model to produce at random, while maintaining responsiveness and call timing from ZF-AIM. Indeed, ZF-AIM-ablated and birds exhibited correlated calling, with the number of calls correlated across bins (*F*_1,1519.2_ = 644.0, *p* < 0.001) and response latencies of ZF-AIM and ZF-AIM-ablated were similar (*F*_1,20_ = 2.4, *p* = 0.134).

When analyzing acoustic flexibility, we observed limited selectivity, modulation, and covariation by ZF-AIM-ablated during interactions with a live bird. Surprisingly, we did observe that the model was selective based on PC1, being more likely to respond to calls that were higher on PC1 (*F*_1,10_ = 19.4, *p* = 0.001; Fig. S5B). Because duration loads strongly on PC1, this results from the model being more likely to produce a call within the 1 second response window when responding to longer calls from the bird (this model begins processing the wait time upon detecting the onset of a partner call). The magnitude of this effect was smaller than observed in live birds. There was no significant selectivity based on PC2 (*F*_1,10_ = 1.6, *p* = 0.241; Fig. S5B). The model exhibited no modulation on PC1, albeit slight modulation on PC2, with responses being lower on PC2 than other calls (PC1: *F*_1,10_ = 1.1, *p* = 0.310; PC2: *F*_1,10_ = 7.5, *p* = 0.021; Fig. S5C). The model exhibited no significant response covariation (PC1: *t*_10_ = −1.9, *p* = 0.080; PC2: *F*_10_ = 1.8, *p* = 0.104; Fig. S5D).

**Figure S5:**
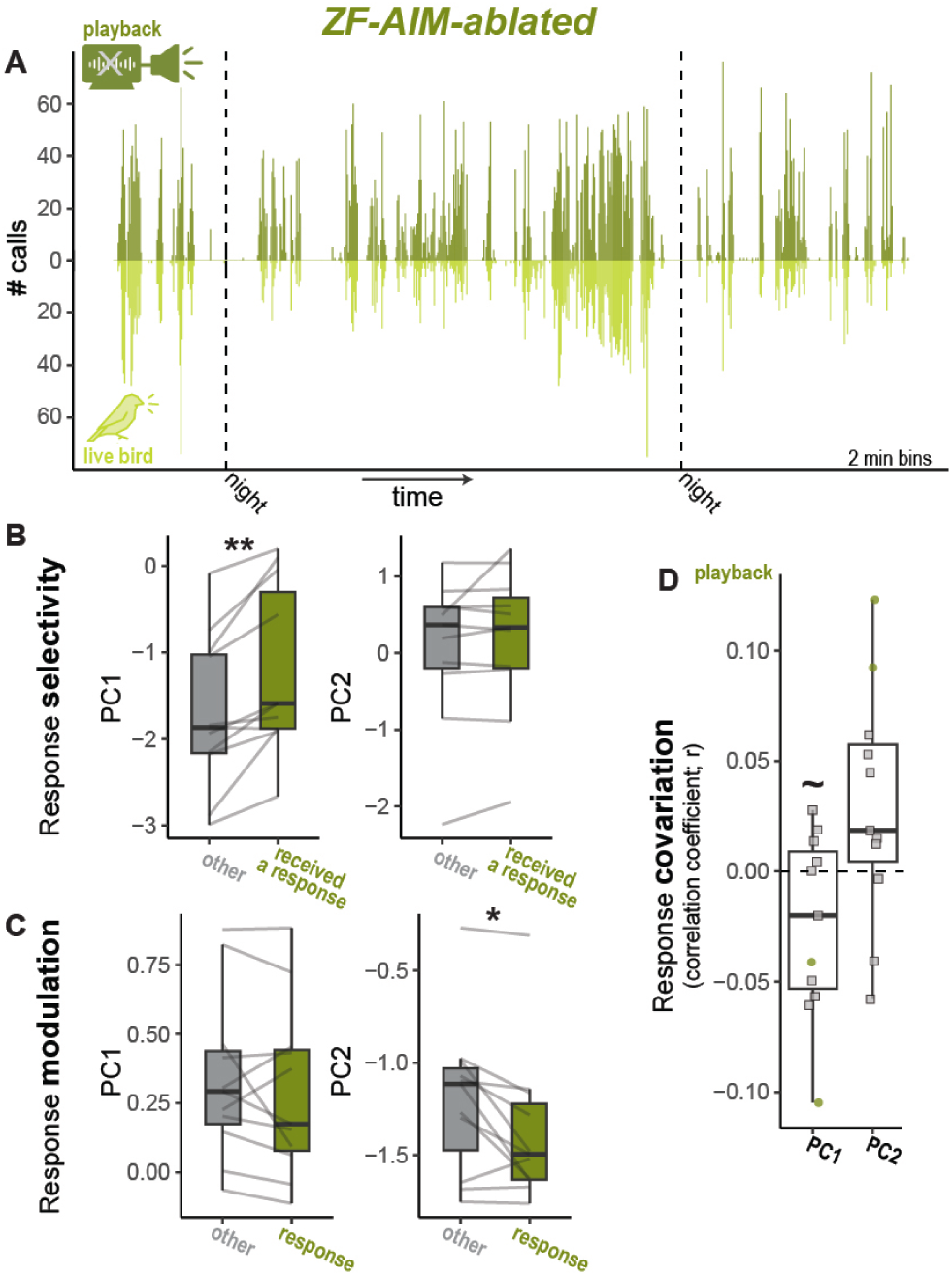
Behavior of ZF-AIM-ablated during live interactions with a bird. A. An example histogram showing the calling behavior of the playback (going up) and the bird (going down) across 2 minute bins. Dashed vertical lines indicate night, when birds do not call and the playback was turned off. B. Response selectivity of the model based on the calls from the live bird for PC1 (left) and PC2 (right). C. Response modulation from ZF-AIM-ablated across its responses to the bird for PC1 (left) and PC2 (right). D. Covariation of acoustic features from ZF-AIM-ablated toward those of the live bird. For all panels ∼ *p* < 0.10, ∗ *p* < 0.05, ∗∗ *p* < 0.005.

**Table S7:** Statistical results from the interactions of live pairs of birds.

| model | Dep | Ind | NumDF | DenDF | stat | p | notes |
| --- | --- | --- | --- | --- | --- | --- | --- |
| LMM | partner calls | self calls | 1 | 5860.3 | F = 3679.5 | < 0.001 |  |
| LMM | total calls | time bin | 1 | 167.5 | F = 25.0 | < 0.001 |  |
| LMM | response rate | time bin | 1 | 481.8 | F = 5.5 | 0.019 |  |
| LMM | PC1 | recieves response | 1 | 92.2 | F = 69.6 | < 0.001 |  |
| LMM | PC2 | recieves response | 1 | 87.3 | F = 5.1 | 0.027 |  |
| LMM | PC1 | is a response | 1 | 89.0 | F = 87.0 | < 0.001 |  |
| LMM | PC2 | is a response | 1 | 95.9 | F = 8.4 | 0.005 |  |
| LMM | PC1 r | NA | 19.4 | NA | t = 15.3 | < 0.001 | intercept only |
| LMM | PC2 r | NA | 17.1 | NA | t = 3.9 | 0.001 | intercept only |
| LMM | r | PC | 1 | 110.6 | F = 395.3 | < 0.001 | PC1 > PC2 |

**Table S8:** Statistical results from passive playbacks.

| model | Dep | Ind | NumDF | DenDF | stat | p | notes |
| --- | --- | --- | --- | --- | --- | --- | --- |
| LMM | mean calls | On vs Off | 1 | 13.0 | F = 70.9 | < 0.001 | fixed |
| LMM | mean calls | On vs Off | 1 | 15.0 | F = 34.2 | < 0.001 | random |
| LMM | Response latency | condition | 2 | 21.5 | F = 4.8 | 0.019 |  |
| post-hoc |  |  | 21 |  | t = -3.1 | 0.014 | natural - fixed |
| post-hoc |  |  | 21 |  | t = -1.4 | 0.345 | natural - random |
| post-hoc |  |  | 22 |  | t = 1.6 | 0.263 | fixed - random |
| LMM | response rate | time | 1 | 667.5 | F = 16.2 | < 0.001 |  |
|  |  | condition | 2 | 63.5 | F = 29.7 | < 0.001 |  |
|  |  | time * condition | 2 | 667.5 | F = 2.5 | 0.085 |  |
| LMM | PC1 Diff | condition | 2 | 20.6 | F = 2.2 | 0.133 | selectivity |
| LMM | PC1 Diff |  | 23.3 |  | t = 6.5 | < 0.001 | selectivity, intercept only: live |
| LMM | PC1 Diff |  | 27.6 |  | t = 3.8 | < 0.001 | selectivity, intercept only: fixed |
| LMM | PC1 Diff |  | 26.1 |  | t = 5.4 | < 0.001 | selectivity, intercept only: random |
| LMM | PC2 Diff | condition | 2 | 34.0 | F = 0.6 | 0.553 | selectivity |
| LMM | PC2 Diff |  | 34.0 |  | t = -3.1 | 0.004 | selectivity, intercept only: live |
| LMM | PC2 Diff |  | 34.0 |  | t = -1.6 | 0.13 | selectivity, intercept only: fixed |
| LMM | PC2 Diff |  | 34.0 |  | t = -1.6 | 0.118 | selectivity, intercept only: random |
| LMM | PC1 Diff | condition | 2 | 34.0 | F = 6.9 | 0.003 | modulation |
| post-hoc |  |  | 23 |  | t = 3.3 | 0.008 | natural - fixed |
| post-hoc |  |  | 23 |  | t = 3.0 | 0.019 | natural - random |
| post-hoc |  |  | 25 |  | t = -0.4 | 0.908 | fixed - random |
| LMM | PC1 Diff |  | 34.0 |  | t = 6.9 | < 0.001 | modulation, intercept only: live |
| LMM | PC1 Diff |  | 34.0 |  | t = 1.7 | 0.102 | modulation, intercept only: fixed |
| LMM | PC1 Diff |  | 34.0 |  | t = 2.4 | 0.024 | modulation, intercept only: random |
| LMM | PC2 Diff | condition | 2 | 21.3 | F = 7.8 | 0.003 | modulation |
| post-hoc |  |  | 21 |  | t = -3.5 | 0.006 | natural - fixed |
| post-hoc |  |  | 21 |  | t = -3.2 | 0.012 | natural - random |
| post-hoc |  |  | 22 |  | t = 0.4 | 0.927 | fixed - random |
| LMM | PC2 Diff |  | 16.6 |  | t = 1.4 | 0.176 | modulation, intercept only: live |
| LMM | PC2 Diff |  | 18.9 |  | t = 3.7 | 0.002 | modulation, intercept only: fixed |
| LMM | PC2 Diff |  | 18.0 |  | t = 3.5 | 0.003 | modulation, intercept only: random |
| LMM | PC1 r | condition | 2 | 23.6 | F = 29.3 | < 0.001 |  |
| post-hoc |  |  | 23 |  | t = 6.7 | < 0.001 | natural - fixed |
| post-hoc |  |  | 22 |  | t = 6.2 | < 0.001 | natural - random |
| post-hoc |  |  | 24 |  | t = -0.6 | 0.806 | fixed - random |
| LMM | PC1 r |  | 32.4 |  | t = 12.9 | < 0.001 | intercept only: live |
| LMM | PC1 r |  | 33.4 |  | t = 3.2 | 0.003 | intercept only: fixed |
| LMM | PC1 r |  | 33.1 |  | t = 4.2 | < 0.001 | intercept only: random |
| LMM | PC2 r | condition | 2 | 20.1 | F = 1.3 | 0.287 |  |
| LMM | PC2 r |  | 29.7 |  | t = 2.6 | 0.013 | intercept only: live |
| LMM | PC2 r |  | 32.3 |  | t = 0.6 | 0.554 | intercept only: fixed |
| LMM | PC2 r |  | 31.5 |  | t = 1.0 | 0.348 | intercept only: random |

**Table S9:** Statistical results from interactions between a live bird and ZF-AIM. Note: “caller” indicates a categorical variable with “bird” or “model” as the two levels.

| model | Dep | Ind | NumDF | DenDF | stat | p | notes |
| --- | --- | --- | --- | --- | --- | --- | --- |
| LMM | bird calls | model calls | 1 | 1433.4 | F = 1218.2 | < 0.001 |  |
| LMM | mean calls | time | 1 | 129.1 | F = 43.2 | < 0.001 |  |
|  |  | caller | 1 | 17.8 | F = 0.4 | 0.519 |  |
|  |  | time * caller | 1 | 129.1 | F = 6.2 | 0.014 |  |
| LMM | response rate | time | 1 | 125.3 | F = 25.3 | < 0.001 |  |
|  |  | caller | 1 | 31.4 | F = 1.2 | 0.284 |  |
|  |  | time * caller | 1 | 125.3 | F = 18.4 | < 0.001 |  |
| LM | PC1 diff | caller | 1 | 20 | F = 0.0 | 0.85 | Selectivity |
| LM | PC1 diff |  | 20 |  | t = 3.6 | 0.002 | intercept only: birds |
| LM | PC1 diff |  | 20 |  | t = 3.8 | 0.001 | intercept only: model |
| LM | PC2 diff | caller | 1 | 20 | F = 15.2 | < 0.001 | Selectivity |
| LM | PC2 diff |  | 20 |  | t = -6.1 | < 0.001 | intercept only: birds |
| LM | PC2 diff |  | 20 |  | t = -0.5 | 0.596 | intercept only: model |
| LM | PC1 diff | caller | 1 | 20 | F = 6.3 | 0.021 | Modulation |
| LM | PC1 diff |  | 20 |  | t = 3.8 | 0.001 | intercept only: birds |
| LM | PC1 diff |  | 20 |  | t = 0.3 | 0.799 | intercept only: model |
| LM | PC2 diff | caller | 1 | 20 | F = 8.5 | 0.009 | Modulation |
| LM | PC2 diff |  | 20 |  | t = 1.8 | 0.084 | intercept only: birds |
| LM | PC2 diff |  | 20 |  | t = -2.3 | 0.033 | intercept only: model |
| LM | PC1 r | caller | 1 | 20 | F = 2.2 | 0.151 |  |
| LM | PC1 r |  | 20 |  | t = 3.9 | < 0.001 | intercept only: birds |
| LM | PC1 r |  | 20 |  | t = 1.8 | 0.084 | intercept only: model |
| LM | PC2 r | caller | 1 | 20 | F = 1.1 | 0.297 |  |
| LM | PC2 r |  | 20 |  | t = 4.5 | < 0.001 | intercept only: birds |
| LM | PC2 r |  | 20 |  | t = 3.0 | 0.007 | intercept only: model |

**Table S10:** Statistical results from comparisons of birds’ behavior across live interactions, interactions with ZF-AIM, and interactions with ZF-AIM-ablated.

| model | Dep | Ind | NumDF | DenDF | stat | p | notes |
| --- | --- | --- | --- | --- | --- | --- | --- |
| LMM | response rate | time | 1 | 395.8 | F = 71.4 | < 0.001 |  |
|  |  | condition | 2 | 65.6 | F = 11.7 | < 0.001 |  |
|  |  | time * condition | 2 | 395.8 | F = 15.7 | < 0.001 |  |
| LMM | PC1 | condition | 2 | 20.0 | F = 1.5 | 0.256 | Selectivity |
| LMM | PC1 |  | 21.3 |  | t = 4.8 | < 0.001 | intercept only: live |
| LMM | PC1 |  | 21.3 |  | t = 4.4 | < 0.001 | intercept only: model |
| LMM | PC1 |  | 21.3 |  | t = 3.1 | 0.005 | intercept only: ablated model |
| LMM | PC2 | condition | 2 | 20.0 | F = 10.4 | < 0.001 | Selectivity |
| post-hoc |  |  | 20 |  | t = 4.6 | < 0.001 | natural - model |
| post-hoc |  |  | 20 |  | t = 2.4 | 0.062 | natural - ablated model |
| post-hoc |  |  | 20 |  | t = -2.1 | 0.109 | model - ablated model |
| LMM | PC2 |  | 23.5 |  | t = -2.1 | 0.044 | intercept only: live |
| LMM | PC2 |  | 23.5 |  | t = -7.2 | < 0.001 | intercept only: model |
| LMM | PC2 |  | 23.5 |  | t = -4.8 | < 0.001 | intercept only: ablated model |
| LMM | PC1 | condition | 2 | 20.0 | F = 0.8 | 0.472 | Modulation |
| LMM | PC1 |  | 20.0 |  | t = 4.4 | < 0.001 | intercept only: live |
| LMM | PC1 |  | 20.0 |  | t = 3.2 | 0.004 | intercept only: model |
| LMM | PC1 |  | 20.0 |  | t = 3.4 | 0.003 | intercept only: ablated model |
| LMM | PC2 | condition | 2 | 20.0 | F = 2.4 | 0.114 | Modulation |
| LMM | PC2 |  | 14.7 |  | t = 1.2 | 0.255 | intercept only: live |
| LMM | PC2 |  | 14.7 |  | t = 1.5 | 0.144 | intercept only: model |
| LMM | PC2 |  | 14.7 |  | t = 2.8 | 0.015 | intercept only: ablated model |
| LMM | PC1 r | condition | 2 | 20.0 | F = 13.0 | < 0.001 |  |
| post-hoc |  |  | 20 |  | t = 4.0 | 0.002 | natural - model |
| post-hoc |  |  | 20 |  | t = 4.8 | < 0.001 | natural - ablated model |
| post-hoc |  |  | 20 |  | t = 0.8 | 0.702 | model - ablated model |
| LMM | PC1 r |  | 26.6 |  | t = 9.3 | < 0.001 | intercept only: live |
| LMM | PC1 r |  | 26.6 |  | t = 4.5 | < 0.001 | intercept only: model |
| LMM | PC1 r |  | 26.6 |  | t = 3.5 | 0.002 | intercept only: ablated model |
| LMM | PC2 r | condition | 2 | 20.0 | F = 4.3 | 0.028 |  |
| post-hoc |  |  | 20 |  | t = -0.6 | 0.835 | natural - model |
| post-hoc |  |  | 20 |  | t = 2.2 | 0.095 | natural - ablated model |
| post-hoc |  |  | 20 |  | t = 2.8 | 0.03 | model - ablated model |
| LMM | PC2 r |  | 29.3 |  | t = 3.6 | 0.001 | intercept only: live |
| LMM | PC2 r |  | 29.3 |  | t = 4.3 | < 0.001 | intercept only: model |
| LMM | PC2 r |  | 29.3 |  | t = 0.6 | 0.522 | intercept only: ablated model |

## Footnotes

1 http://www.audacityteam.org/

2 https://www.ravensoundsoftware.com/software/raven-pro/

3 Implementation from https://github.com/lucidrains/recurrent-memory-transformer-pytorch.

4 Implementation from https://github.com/facebookresearch/audiocraft

5 https://github.com/facebookresearch/audiocraft/blob/main/config/model/encodec/encodec_base_causal.yaml

6 https://github.com/facebookresearch/audiocraft/blob/main/config/solver/compression/default.yaml

7 https://github.com/facebookresearch/audiocraft/blob/main/config/model/encodec/encodec_base_causal.yaml

8 https://github.com/facebookresearch/audiocraft/blob/main/config/solver/compression/default.yaml

9 We use the FAD implementation from https://github.com/gudgud96/frechet-audio-distance

10 https://github.com/spatialaudio/python-sounddevice/

11 https://osf.io/wf2x8/overview?view_only=7d7b699afec949a089c8b7abfd8e71c2

12 https://osf.io/wf2x8/overview?view_only=7d7b699afec949a089c8b7abfd8e71c2

13 https://osf.io/wf2x8/overview?view_only=7d7b699afec949a089c8b7abfd8e71c2

